# AKIR-1 Regulates Proteasome Localization and Function in *Caenorhabditis elegans*

**DOI:** 10.1101/2022.10.03.510264

**Authors:** Johanna Pispa, Elisa Mikkonen, Leena Arpalahti, Congyu Jin, Carmen Martínez-Fernández, Julián Cerón, Carina I. Holmberg

## Abstract

Regulated protein clearance is vital for cells to maintain protein homeostasis and the conditions essential for survival. The primary machinery for intracellular protein degradation is the ubiquitin– proteasome system (UPS), by which ubiquitin-tagged proteins are degraded by the proteasome. Proteasomes are present both in the cytoplasm and the nucleus, but the mechanisms coordinating proteasome activity and its subcellular localization in a multicellular organism are still unclear. Here, we identified the nuclear protein-encoding gene *akir-1* as a proteasome regulator in a genome-wide *Caenorhabditis elegans* (*C. elegans*) RNAi screen. We show that the depletion of *akir-1* causes accumulation of endogenous polyubiquitinated proteins in the nuclei of intestinal cells, concomitant with slower *in vivo* proteasomal degradation in this subcellular compartment. Remarkably, the loss of *akir-1* does not induce an accumulation of polyubiquitinated proteins in oocyte nuclei, though *akir-1* is essential for the nuclear localization of proteasomes in both cell types. We further show that the importin family member *ima-3* genetically interacts with *akir-1*, and affects subcellular distribution of polyubiquitinated proteins in intestinal cells. We show for the first time that conserved AKIR-1 is important for the nuclear transport of proteasomes in a multicellular organism, suggesting a role for AKIR-1 in the maintenance of proteostasis.

## Introduction

Protein degradation is one of the essential mechanisms that the cell uses to maintain its homeostasis and protein balance. The major machinery for degrading individual proteins is the ubiquitin– proteasome system (UPS), by which a cascade of enzymes for ubiquitin activation (E1), conjugation (E2), and ligation (E3) marks selected proteins with polyubiquitin chains as a signal for degradation (Kleiger & Mayor, 2014). Hydrolysis of the tagged proteins into small peptide fragments is performed by the proteasome, a large 2,5 MDa protein complex composed of two subcomplexes: a catalytic core particle (CP or 20S) and a regulatory particle (RP or 19S). The catalytic activities, i.e., trypsin-, chymotrypsin- and caspase-like activity of the barrel-shaped 20S particle, are contained within the two seven-subunit β-rings, which are stacked between two seven-subunit α-rings. The core particle can be capped at either one (26S) or both ends by the multisubunit regulatory particle, which functions in recognition, binding, deubiquitination, unfolding, and translocation of the substrates into the lumen of the core particle (Sakata et al., 2021). The proteasomal degradation capabilities are broadened with the ability of the free core particle to degrade some proteins independently of ubiquitination (Demasi & da Cunha, 2018) and the existence of proteasome variants with alternative subunits of the core particle or the regulatory particle, or activating complexes (Budenholzer et al., 2017).

During the last decades, studies based on biochemical and immunological methods and live cell imaging have reported proteasomes to be localized both in the cytoplasm and in the nucleus in various cell types (Wojcik & DeMartino, 2003). Despite vast accumulating evidence on variation in nuclear and cytoplasmic distribution of proteasomes in different cell lines and physiological conditions, the regulatory mechanisms governing nuclear localization and function of the proteasomes are only beginning to emerge (Enenkel et al., 2022). Generally, controlled UPS-mediated nuclear degradation is important for the regulation of many nuclear proteins, such as several transcription factors, histones or epigenetic modifying enzymes (von Mikecz, 2006; Geng et al., 2012; Bach & Hegde, 2016). However, there are also reports showing that both in yeast and human cells nuclei lack proteolytic activity, suggesting that proteasomes localize mainly on the cytoplasmic side of the nuclear membrane (Enenkel et al., 1998; Dang et al., 2016). In addition, in some circumstances ubiquitinated proteins seem to be exported from the nucleus for cytoplasmic degradation in the cytoplasm (Hirayama et al., 2018).

In dividing cells, proteasomes can enter the nucleus by simple diffusion during nuclear membrane breakage. However, in non-dividing cells or in yeast with closed mitosis, proteasomes or their subcomplexes are transported via the nuclear pores (Savulescu et al., 2011; Pack et al., 2014). In yeast, import of proteasomes to the nucleus requires the importin α family member SRP1 (Lehmann et al., 2002). Recently, studies have further shown that importin 9 is involved in translocation of proteasomes into the nucleus in human cancer cell lines, and during *Drosophila melanogaster (D. melanogaster)* spermatogenesis (de Almeida et al., 2021; Palacios et al., 2021). In *Saccharomyces cerevisiae (S. cerevisiae) and Schizosaccharomyces pombe (S. pombe)*, the adapter proteins Sts1 and Cut8 respectively bind proteasome and mediate nuclear import of proteasomes (Tatebe & Yanagida, 2000; Takeda & Yanagida, 2005; Chen et al., 2011; Takeda et al., 2011; Budenholzer et al., 2020). Additionally, the alternative regulatory protein Blm10 has been identified as an adapter in nuclear import of core particles in yeast (Weberruss et al., 2013).

We have previously demonstrated that proteasomes are localized both in the cytoplasm and in the nucleus in multiple tissues in *C. elegans* (Mikkonen et al., 2017). Here, we show that the conserved nuclear protein–encoding gene *akir-1*, which we identified in a genome-wide RNAi screen for proteasome regulators, is required for nuclear localization of proteasomes in *C. elegans*, and that depletion of *akir-1* causes nuclear accumulation of polyubiquitinated proteins in a tissue-specific manner. In addition, downregulation of the *C. elegans* importins *ima-3* and *imb-1* mimics the *akir-1* RNAi–induced polyubiquitin phenotype in intestinal cells. Our results are the first report showing the importance of AKIR-1 in proteasome nuclear transport and protein homeostasis in a multicellular organism.

## Results

### Loss of akir-1 results in nuclear accumulation of polyubiquitinated proteins in C. elegans intestinal cells

We identified *akir-1* as one hit gene, out of more than 30 potential new regulators of the proteasome, in a genome-wide RNAi screen (screen data not included here), which we performed on a transgenic *C. elegans* strain expressing the polyubiquitin-binding fluorescent reporter in intestinal cells (Matilainen et al., 2013). Increased fluorescence reflects stabilization of the reporter upon accumulation of endogenous polyubiquitinated proteins, and we have previously shown that the reporter responds to changes in proteasome activity and physiological stimuli (Matilainen et al., 2013; Jha & Holmberg, 2020; Pispa et al., 2020). We discovered that downregulation of *akir-1* (*E01A2*.*6*), increased the reporter fluorescence (mean fold induction 4.5 +/- SD 2.5, n = 3 independent experiments), and in addition, strikingly changed its uniform and diffuse cellular distribution into a more converged pattern (Figure 1A). Hoechst staining of DNA confirmed that the new fluorescence pattern resulted from the polyubiquitin-binding reporter concentrating mainly into the intestinal nuclei, excluding the nucleoli (Figure 1B), of *C. elegans*.

**Figure 1.**
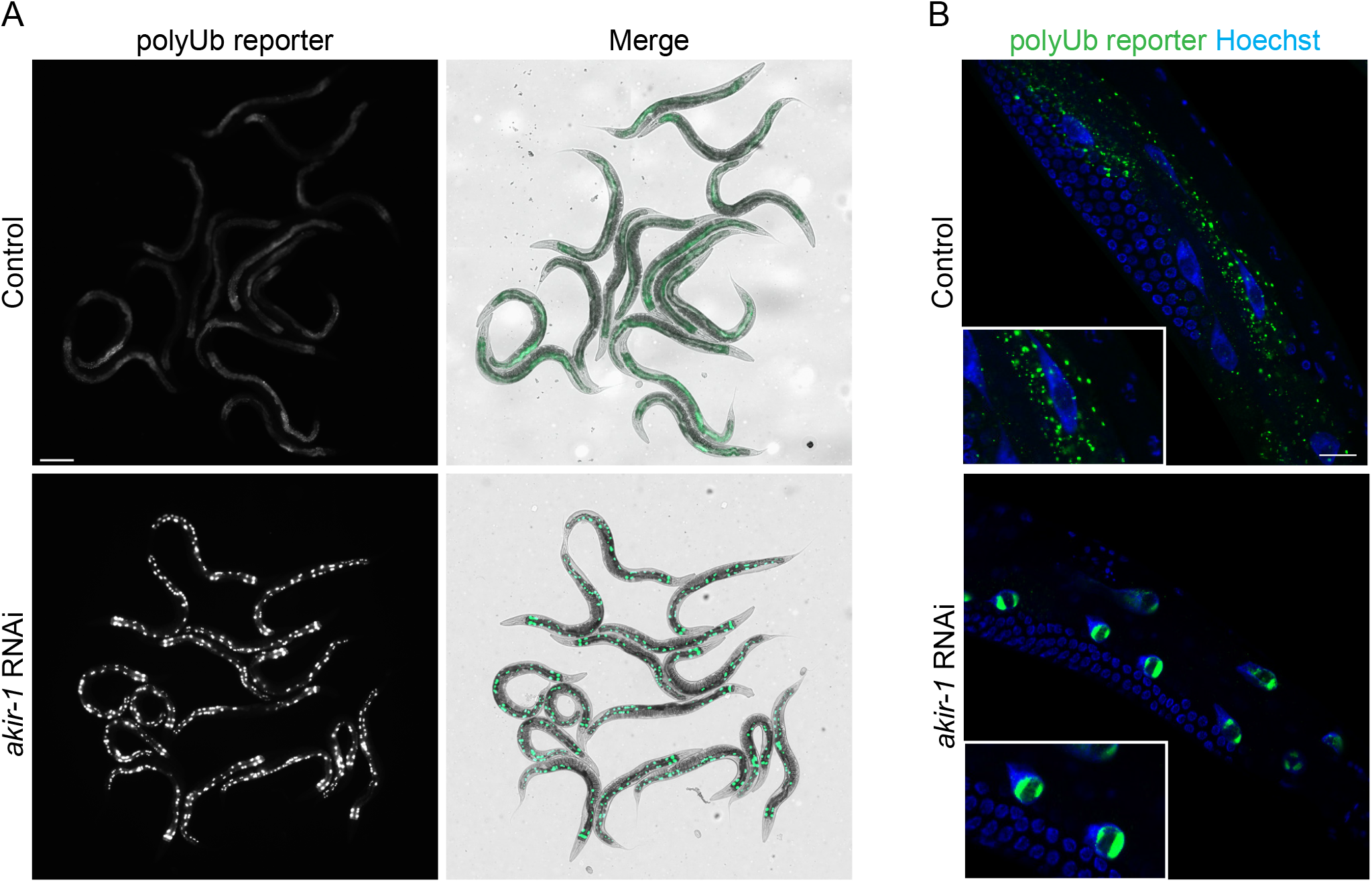
The polyubiquitin-binding reporter accumulates in the intestinal nuclei upon *akir-1* RNAi. (**A**) Representative fluorescence micrographs of control and *akir-1* RNAi-treated N2 animals expressing the polyubiquitin (polyUb) reporter (*vha-6p::UIM2-ZsProSensor*) in the intestinal cells (left panels). Merge represents overlay of fluorescence and bright-field images (right panels). Scale bar, 200 pm. (**B**) Representative confocal micrographs of control and akir-1 RNAi-treated polyubiquitin reporter animals with Hoechst-visualized nuclei. Insets show enlargements. Scale bar, 20 µm.

Our results with the polyubiquitin-binding reporter strain suggest that *akir-1* downregulation causes accumulation of endogenous polyubiquitinated proteins in intestinal nuclei. We next used an anti-polyubiquitin antibody for immunostaining of dissected intestines from wild-type N2 animals treated with control or *akir-1* RNAi. We used dissected tissues to improve antibody tissue penetrance during immunostaining. Although both control and *akir-1* RNAi–treated animals showed positive nuclear staining of polyubiquitinated proteins in the intestine, the portion of animals with positive nuclear staining clearly increased from 60% to 90% upon *akir-1* RNAi (Figure 2A). Importantly, when the accumulating polyubiquitin immunostaining in the intestine nuclei was visually estimated as none (i.e., less or equal to cytoplasmic immunostaining), weak or strong, the number of animals with strong immunostaining increased from 14% in control to 72% upon *akir-1* RNAi treatment (Figure 2A). As qPCR analysis showed that *akir-1* mRNA was downregulated efficiently, but not completely, after *akir-1* RNAi treatment (Figure 2 ─ figure supplement 1A), we examined polyubiquitinated proteins also in the *akir-1(gk528)* deletion mutant, which is predicted to be a null allele (Clemons et al., 2013). In accordance with the *akir-1* RNAi results, *akir-1* mutants displayed a clear increase in nuclear polyubiquitin immunostaining compared to wild-type animals (90% *akir-1(gk528)*, 44% control), and in particular, the strong nuclear immunostaining increased from 16% in wild-type animals to 74% in *akir-1* mutants (Figure 2B). To investigate the total levels of endogenous polyubiquitinated proteins, we performed Western blot analysis of whole animal lysates with an anti-polyubiquitin antibody. Variability among independent experiments was observed, but no systematic difference in the amount of polyubiquitinated proteins was detected in lysates of *akir-1* mutants compared to control lysates (Figure 2 ─ figure supplement 1B). Taken together, our results show that depletion of *akir-1* leads to accumulation of polyubiquitinated proteins in the nucleus of intestinal cells in *C. elegans*.

**Figure 2.**
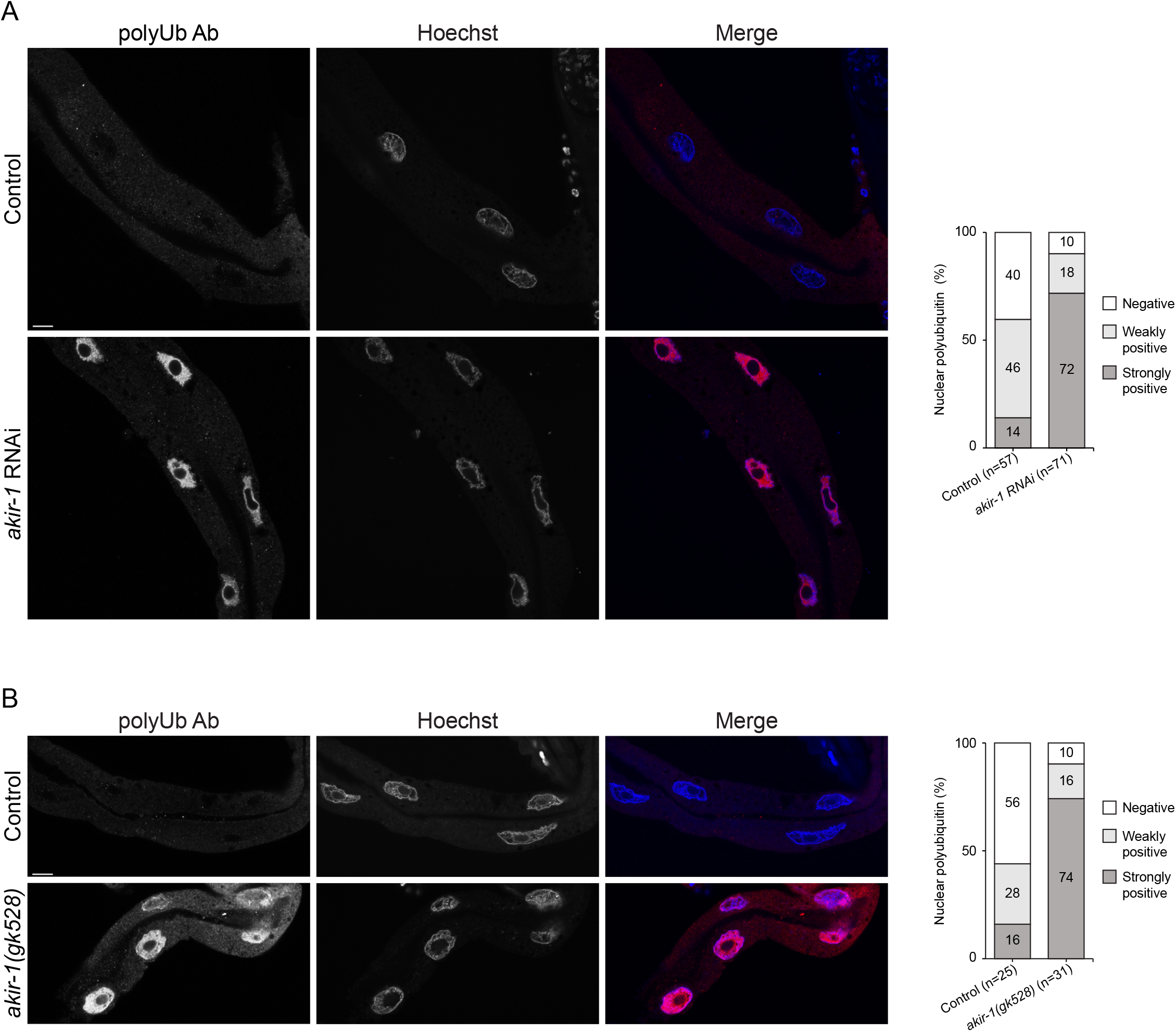
Depletion of *akir-1* leads to nuclear accumulation of immunostained polyubiquitinated proteins in the intestine. (**A, B**) Representative confocal micrographs of polyubiquitin immunostaining (polyUb Ab) in dissected intestines of control and *akir-1* RNAi-treated wild-type (N2) animals (**A**), and in control (N2) animals and akir-1 (*gk528*) mutants (**B**). Nuclei are visualized with Hoechst. Scale bars, 10 pm. The graphs (on right) show visually quantified nuclear polyubiquitin accumulation in the intestine in control and *akir-1* RNAi-treated wild-type animals (**A**), and in control (N2) and *akir-1* (*gk528*) animals (**B**). n = total number of animals. Proportions of animals with strongly positive, weakly positive, or negative (ie., less or equal to cytoplasmic immunostaining) nuclear immunostaining are indicated in percentages.

### Knockdown of akir-1 alters proteasome activity

Accumulation of polyubiquitinated proteins is commonly caused by altered proteasome levels or activity. We first analyzed the total amount of endogenous proteasomes in whole animal lysates by Western blot analysis using an antibody against 20S alpha subunits. No clear change in 20S levels was detected in lysates of either *akir-1* mutants (Figure 3A) or *akir-1* RNAi–treated animals (Figure 3B) compared to control animals. Next, we performed an in-gel proteasome activity assay on whole animal lysates, and detected a slight increase in total proteasome activity upon *akir-1* RNAi (Figure 3C, left graph). Interestingly, the 20S core particle (CP) appears to be the main contributor of this increased activity (Figure 3C). To specifically measure proteasome activity in intestinal cells of *C. elegans*, we employed our previously established UPS reporter animals expressing the photoconvertible proteasomal substrate UbG76V-Dendra2 in the intestine (Hamer et al., 2010; Li et al., 2011). In these animals, a decrease in the amount of photoconverted UbG76V-Dendra2 reporter reflects *in vivo* proteasome activity. We first measured the fluorescence of the photoconverted UbG76V-Dendra2 in the whole intestine, and detected a similar rate of degradation in control and *akir-1* RNAi–treated animals (Figure 4A, extended in Figure 4 ─ figure supplement 1A). As *akir-1* depletion leads to nuclear accumulation of polyubiquitinated proteins, we next monitored photoconverted UbG76V-Dendra2 fluorescence intensity at single-cell level, specifically in the nucleus and cytoplasm, in living animals. The fluorescence of UbG76V-Dendra2 at 18 hours after photoconversion was more intense in the intestinal nuclei than in the cytosol upon *akir-1* RNAi (Figure 4B), demonstrating that *akir-1* knockdown results in slower proteasomal degradation in intestinal nuclei *in vivo*. Taken together, our results suggest that AKIR-1 influences subcellular proteasome activity.

**Figure 3.**
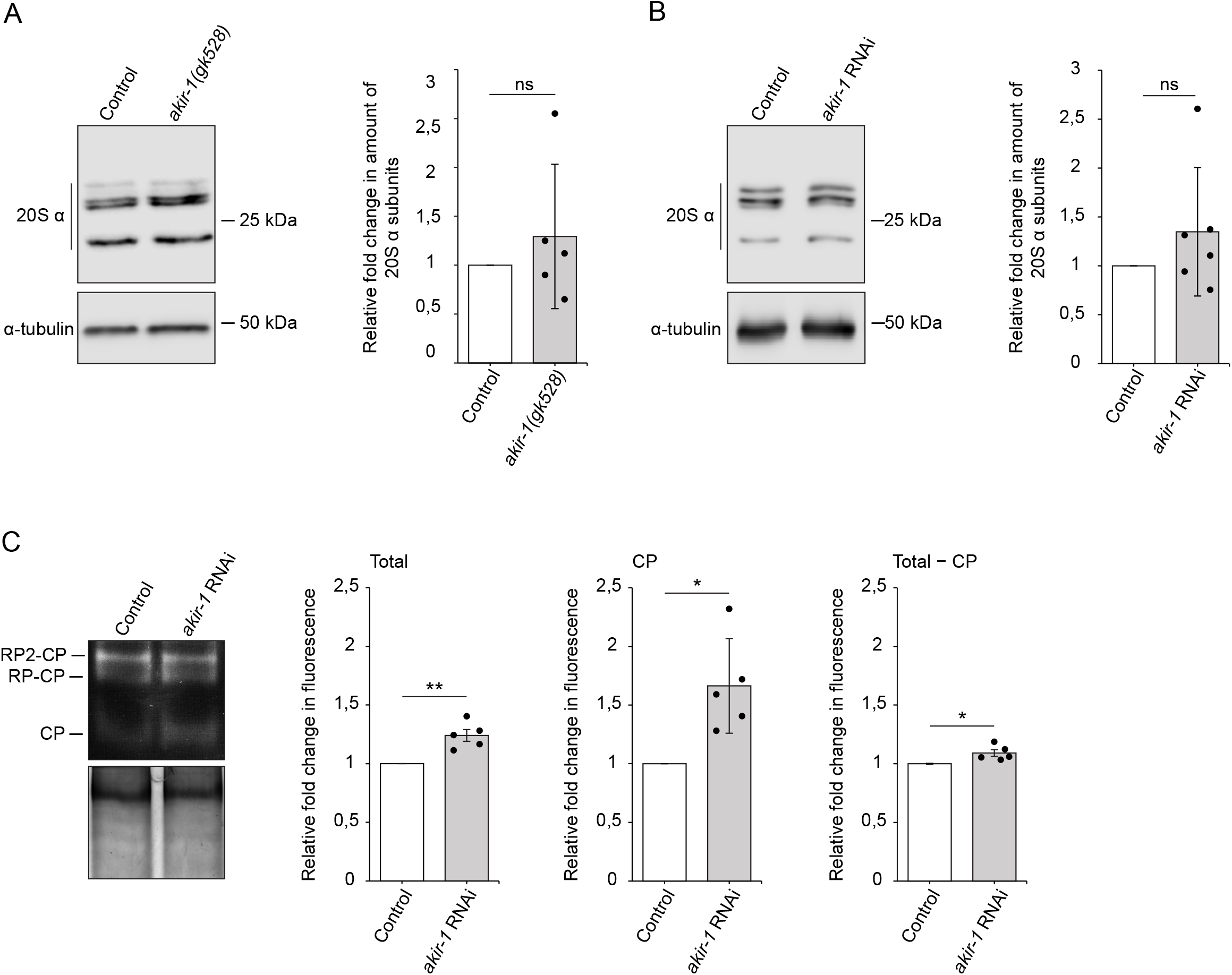
Depletion of *akir-1* does not affect total proteasome amount, but slightly alters *in vitro* proteasome activity. (**A, B**) Western blot analysis with antibody against proteasomal 20S alpha subunits using lysates of *akir-1* (*gk528*) mutants and control (N2) animals (**A**), and of control and *akir-1* RNAi-treated wild-type animals (**B**). Anti-alpha-tubulin antibody was used as a normalization control. The quantification graphs show the average fold change in *akir-1* (*gk528*) mutants compared to control (N2) animals (set to 1; n = 5 independent experiments) (**A**), and in *akir-1* RNAi-treated wild-type animals compared to control RNAi-treated animals (set to 1; n = 6 independent experiments) (**B**). (**C**) In-gel proteasome activity assay with whole animal lysates of wild-type animals exposed to control or *akir-1* RNAi treatment (upper gel). Coomassie staining of the same gel (lower gel). The quantifications show the average fold change in chymotrypsin activity in *akir-1* RNAi-treated wild-type animals compared to control RNAi-treated animals (set to 1; n = 5 independent experiments). Proteasome activity is indicated as total (Total; CP + RP-CP + RP2-CP), as core particle (CP), and as CP activity substracted from total activity (Total - CP). RP, Regulatory particle; Error bars, SD; ns, not significant; *p < 0,05; **p< 0,001.

**Figure 4.**
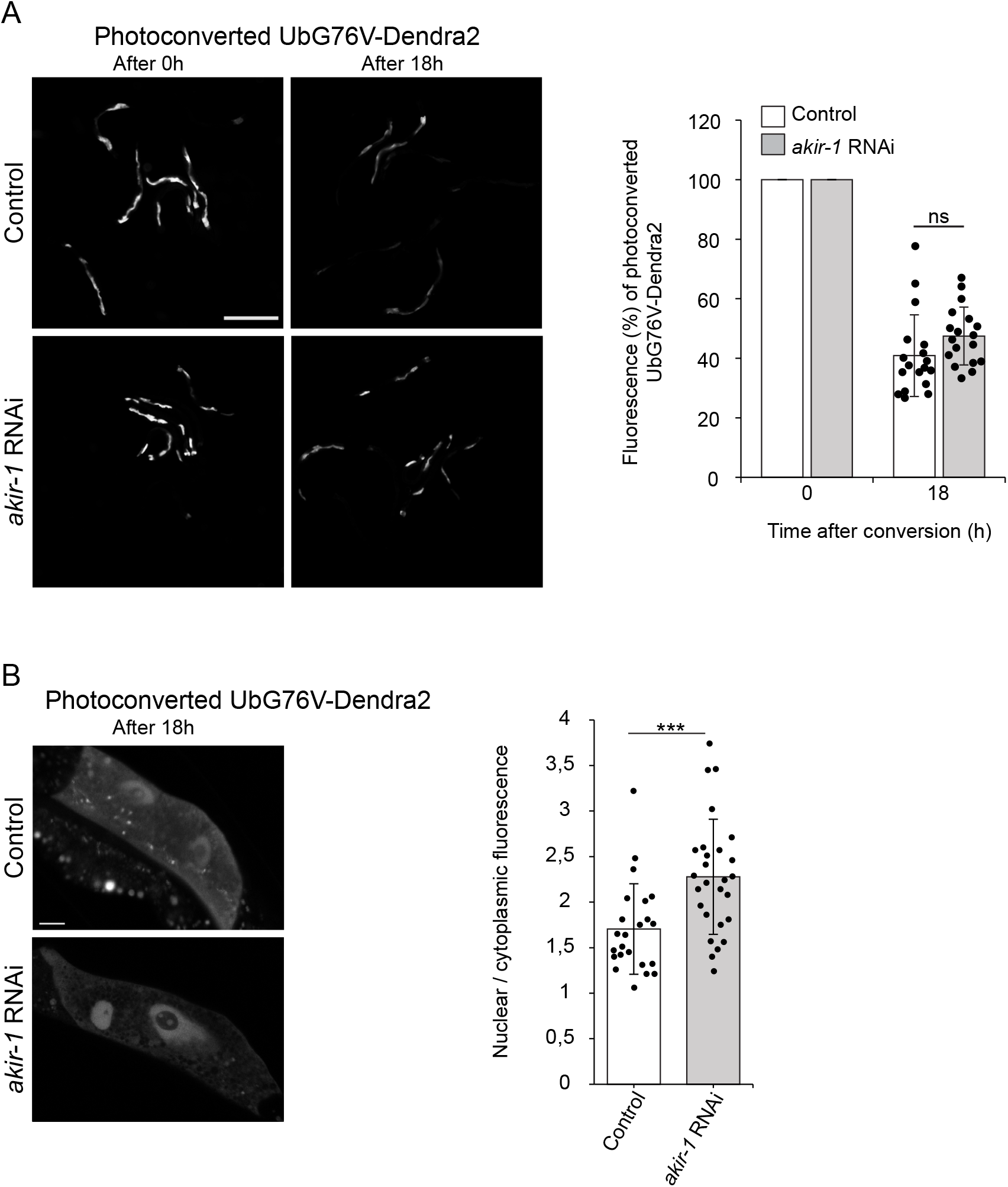
Downregulation of *akir-1* slows degradation of UPS reporter proteins in intestinal nuclei of *C. elegans*. (A) Representative fluorescence micrographs of control and *akir-1* RNAi-treated transgenic *C. elegans* expressing photoconvertible UbG76V-Dendra2 reporter (*vha-6p::UbG76V::Dendra2*) in intestinal cells. The 0 hour (left panels) and 18 hours (right panels) indicate time after photoconversion. Scale bar, 500 μm. The graph shows the mean percentages of fluorescence intensity of the photoconverted UbG76V-Dendra2 18 hours after the photoconversion relative to the fluorescence at the point of photoconversion (0 hour, set as 100%); n = 6 independent experiments with triplicate images of 6-7 animals per image (total number of animals is 108 per treatment); Error bars, SD; ns, not significant. (**B**) Representative confocal fluorescence micrographs of intestinal cells with photoconverted UbG76V-Dendra2 (18 h after conversion) in transgenic *C. elegans* treated with control or *akir-1* RNAi. Scale bar, 10 μm. The graph shows the ratio between nuclear and cytoplasmic mean fluorescence per cell, n = 2 independent experiments (total number of nuclei is 23 in control RNAi and 27 in *akir-1* RNAi treatment); ***p < 0,001; Error bar, SD.

### Intestinal nuclei display reduced proteasome levels upon akir-1 depletion

As the accumulation of polyubiquitinated proteins and the altered *in vivo* proteasome activity in the intestine of *akir-1* mutant and RNAi–treated animals occur specifically in the nucleus, we investigated subcellular distribution of the proteasome by immunostaining dissected intestines with the anti-20S antibody. We have previously shown using this antibody that the proteasome is concentrated in nuclei in the intestine of wild-type *C. elegans* (Mikkonen et al., 2017). Consistently, dissected intestines of wild-type animals displayed high immunofluorescence in the nuclei (Figure 5A and Figure 5 ─ figure supplement 1A). Compared to the controls, both *akir-1* mutants and *akir-1* RNAi– treated animals showed a slight but consistent reduction in 20S immunofluorescence intensity in intestinal nuclei (Figure 5A, Figure 5 ─ figure supplement 1A). As a complementary approach, we investigated the proteasome subcellular localization using transgenic *C. elegans* expressing extrachromosomal arrays of GFP-tagged RPT-5, a subunit of the 19S RP particle of the proteasome, under its endogenous *rpt-5* promoter. The transgenic animals showed a mosaic fluorescence pattern of the ubiquitously expressed GFP::RPT-5. GFP::RPT-5 animals displayed a stronger nuclear fluorescence compared to the cytoplasmic fluorescence in intestinal cells, and upon *akir-1* RNAi a clear reduction in nuclear fluorescence was also observed (Figure 5B), indicating nuclear decrease of 26S proteasome in the intestine. In addition, we investigated the localization of the proteasome-associated deubiquitinase (DUB) UBH-4. UBH-4, and its human homologue UCHL5, have previously been shown to interact with the 19S subunit RPN-13 (Hamazaki et al., 2006; Qiu et al., 2006; Yao et al., 2006; Matilainen et al., 2013) and to be broadly expressed in *C. elegans* tissues, including the intestine (Matilainen et al., 2013). Here, we used a CRISPR-engineered UBH-4::GFP strain (Vicencio et al., 2019), and observed that these animals displayed strong nuclear fluorescence in intestinal nuclei similar to the pattern detected with the anti-20S antibody (Figures 5C and 5A). In accordance with the 20S immunofluorescence and GFP::RPT-5 results, UBH-4::GFP fluorescence decreased in the intestinal nuclei of animals exposed to *akir-1* RNAi when compared to control animals (Figure 5). Together, these results demonstrate that proteasome levels decrease in intestinal nuclei upon the loss of AKIR-1.

**Figure 5.**
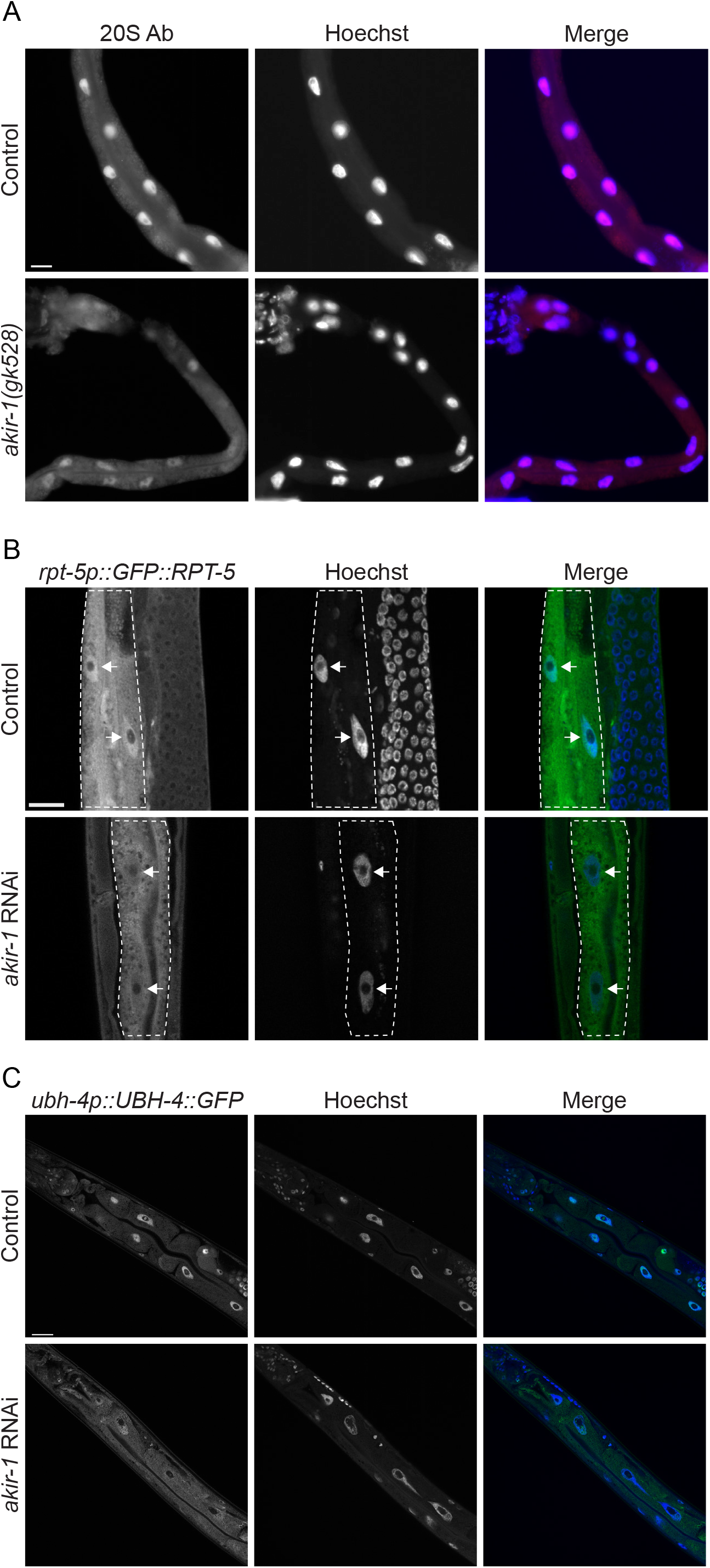
Depletion of *akir-1* decreases proteasome expression in the nuclei of intestinal cells. (**A**) Representative micrographs of proteasome immunostaining (20S Ab) in dissected intestines of control (N2) animals and *akir-1* (*gk528*) mutants. (**B**) Representative confocal micrographs showing GFP fluorescence ratio between nuclei and cytoplasm of control and *akir-1* RNAi-treated *rpt-5p::GFP::RPT-5* animals (40X magnification). Intestinal cells are outlined with white dashed lines and white arrows point to intestinal cell nuclei. (C) Representative confocal micrographs of *ubh-4p::UBH-4::GFP* animals (40X magnification). Nuclei are visualized with Hoechst. Scale bars, 20 µm.

### Depletion of akir-1 affects nuclear polyubiquitinated proteins differently in oocytes and body-wall muscle cells compared to intestinal cells

Whilst performing 20S immunostaining with dissected *akir-1* mutants, we noticed that loss of *akir-1* altered proteasome subcellular distribution not only in the intestinal cells, but also in the oocytes. Oocytes of wild-type animals displayed cytoplasmic and strong nuclear 20S immunofluorescence (Figure 6A, Figure 6 ─ figure supplement 1A). Compared to the wild-type animals, nuclear proteasome localization is clearly reduced in oocytes of *akir-1* mutants (Figure 6A). A similar decrease in intensity of nuclear 20S immunostaining was also detected in oocytes of animals exposed to *akir-1* RNAi (Figure 6 ─ figure supplement 1A). Further, analysis of UBH-4::GFP animals treated with *akir-1* RNAi confirmed reduced levels of nuclear proteasomes in oocytes (Figure 6B). Taken together, these results show that the depletion or downregulation of *akir-1* causes a decrease in nuclear proteasomes in oocytes, similarly to our observation in intestinal cells. However, despite the clear reduction in nuclear proteasomes, we did not observe accumulation of polyubiquitinated proteins in the oocyte nuclei of *akir-1* mutants (Figure 7A). Most oocytes showed a relatively uniform cytoplasmic polyubiquitin immunofluorescence pattern, with weaker signal in the nuclei of both control animals and *akir-1* mutants (Figure 7A). Occasionally, some polyubiquitin-positive staining was detected at the rim of the nuclear membrane in *akir-1* depleted animals (Figure 7A, lower panel). The polyubiquitin staining pattern in oocytes of *akir-1* RNAi–treated animals resembled the results of *akir-1* mutants (Figure 7 ─ figure supplement 1A). Importantly, our results revealed that polyubiquitinated proteins do not accumulate to a similar degree in oocyte nuclei, as in intestinal nuclei, upon loss of *akir-1*.

**Figure 6.**
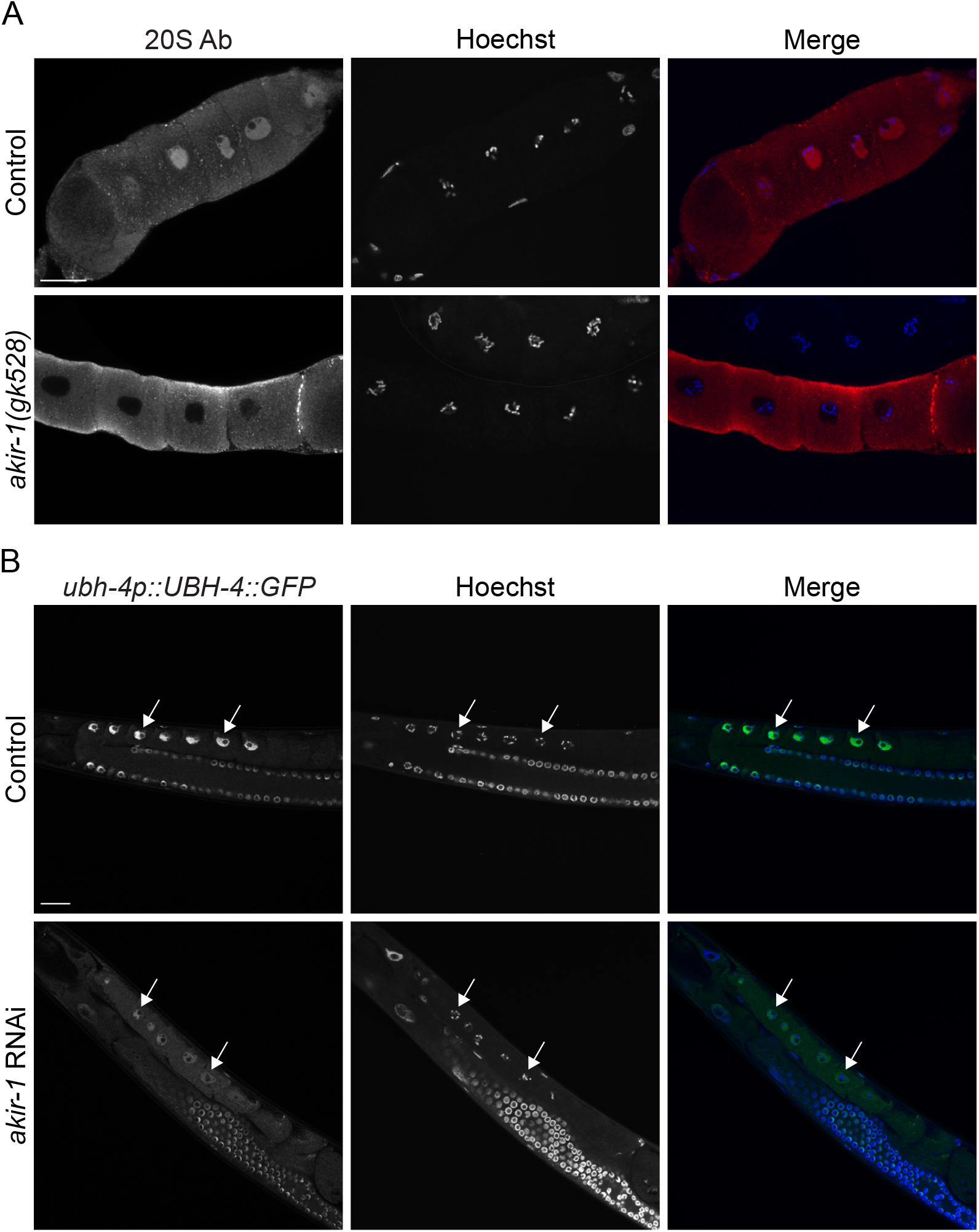
Nuclear proteasome expression decreases in oocytes upon *akir-1* depletion. (**A**) Representative confocal micrographs of proteasome immunostaining (20S Ab) in dissected oocytes of control (N2) animals and *akir-1 (gk528)* mutants. (**B**) Representative confocal micrographs showing GFP fluorescence in oocytes of control and *akir-1* RNAi-treated *ubh-4p::UBH-4::GFP* animals. White arrows point to oocyte nuclei. Nuclei are visualized with Hoechst. Scale bars, 20 µm.

**Figure 7.**
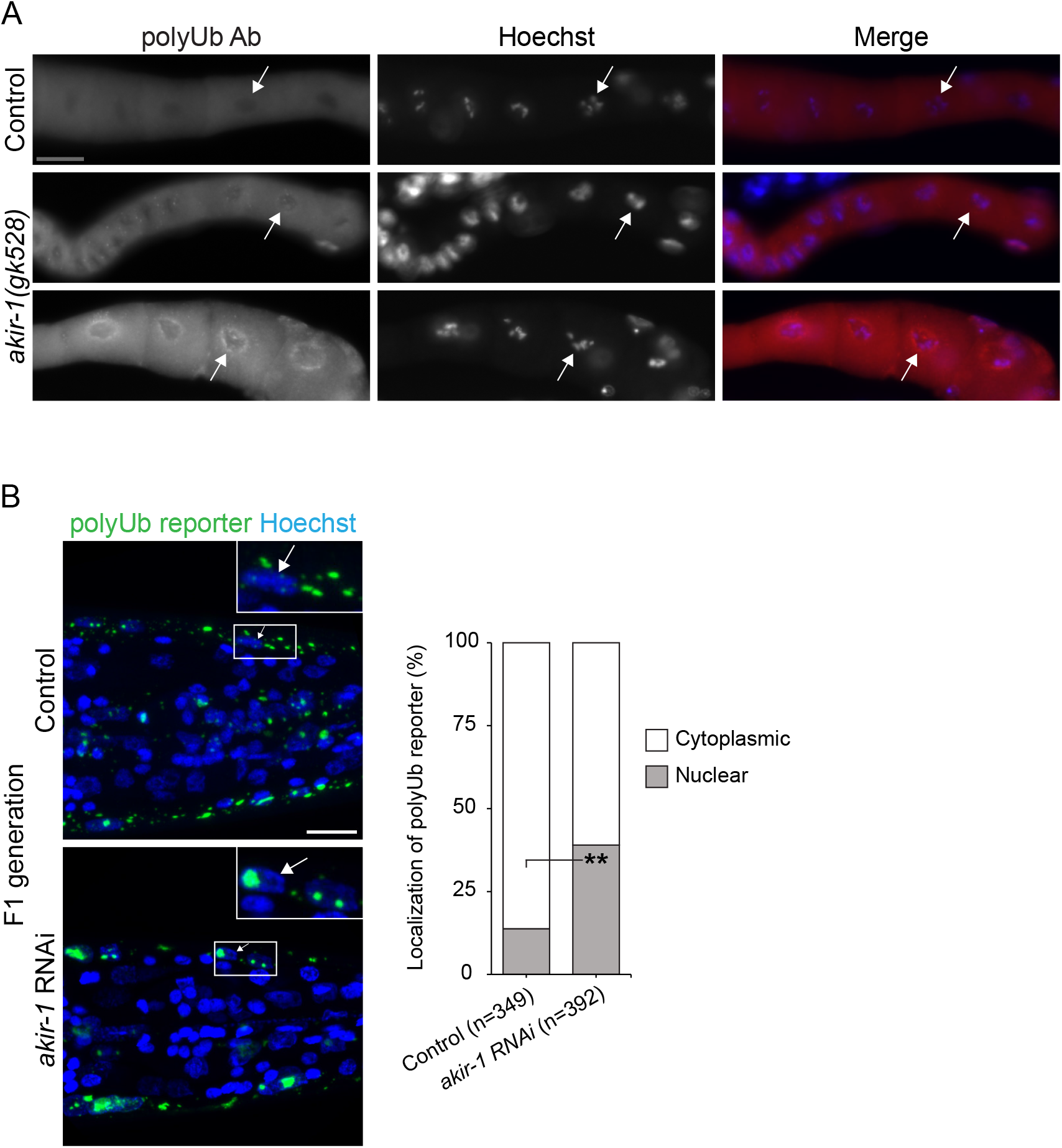
The effects of *akir-1* depletion on nuclear accumulation of polyubiquitinated proteins in oocytes and body-wall muscle cells. (**A**) Representative micrographs of polyubiquitin immunostaining (polyUb Ab) in dissected oocytes of control (N2) animals and *akir-1(gk528)* mutants. Nuclei are visualized with Hoechst. Scale bar, 20 µm. White arrows point to representative nuclei. (**B**) Representative fluorescence micrographs of F1 generation of control and *akir-1* RNAi-treated *rrf-3(pk1426)* animals expressing the polyubiquitin (polyUb) reporter (*unc-54p::UIM2-ZsProSensor)* in the body-wall muscle cells. Nuclei are visualized with Hoechst. Insets show enlargements of the indicated areas. White arrows point to representative nuclei. Scale bar, 10 µm. The graph (on right) shows quantified subcellular localization of polyUb reporter fluorescence in the body-wall muscle cells, n = total number of nuclei; **p < 0,01.

AKIR-1 is required for *C. elegans* muscle development, integrity, and function (Bowman et al., 2020). Therefore, we tested the impact of the loss of *akir-1* on polyubiquitinated proteins in muscles using our previously established transgenic strain expressing the polyubiquitin-binding reporter in the body-wall muscle cells (Jha & Holmberg, 2020; Pispa et al., 2020). Quantification of the mean fluorescent intensity of the polyubiquitin-binding reporter in live animals showed no difference between control and *akir-1* RNAi treatments either in wild-type (N2) background (mean fold change 1.15 +/- SD 0.19, n = 3 independent experiments), or in the RNAi sensitive *rrf-3* background (mean fold change 0.94 +/- SD 0.09, n = 2 independent experiments), and no difference in the fluorescence pattern itself was observed (Figure 7 ─ figure supplement 1B). We also assessed proteasome activity using the photoconvertible UbG76V-Dendra2 reporter expressed in body-wall muscle cells of *C. elegans* (Hamer et al., 2010). No difference in the degradation of photoconverted UbG76V-Dendra2 was detected in *akir-1* RNAi–treated animals compared to control RNAi–treated animals (Figure 7 – figure supplement 1C).

As the RNAi experiments were performed by placing stage 1 larvae (L1) on RNAi plates and assessing the phenotype at day 1 of adulthood, we investigated whether remaining maternal AKIR-1 contribution might influence the lack of polyubiquitin accumulation in body-wall muscle cells. To this end, polyubiquitin-binding reporter animals (in *rrf-3* background) were continuously exposed to *akir-1* RNAi, and their F1 offspring treated in similar manner were monitored at day 1 of adulthood. We assigned reporter fluorescence signals either as nuclear or non-nuclear, based on colocalization with Hoechst nuclear staining. A slight increase in nuclear localization of the reporter was detected in *akir-1* RNAi–treated F1 animals (control 14%, *akir-1* RNAi 39% nuclear signal) (Figure 7B). Taken together, our results suggest that body-wall muscle cells respond differently to *akir-1* depletion in terms of nuclear accumulation of polyubiquitinated proteins than oocytes or intestinal cells.

### Perturbed nuclear transport mimics the akir-1 RNAi-induced polyubiquitin phenotype

AKIR-1 and its metazoan homologues, the Akirin proteins, have been implicated in several physiological processes (Bosch et al., 2020). In *C. elegans*, AKIR-1 is required for meiosis and development (Clemons et al., 2013; Polanowska et al., 2018; Bowman et al., 2020). To investigate phenotypes induced by *akir-1* downregulation under our experimental conditions, we compared the progeny number and lifespan of *rrf-3* animals treated with *akir-1* or control RNAi. We observed a 40% reduction in the number of progeny (Figure 8A) and a 2-day decrease in mean lifespan (Figure 8B, 8C) upon *akir-1* downregulation, which are in an agreement with previous reports on the *akir-1* mutant and an epidermis-specific RNAi strain (Clemons et al., 2013; Polanowska et al., 2018). To exclude that our detected effects on polyubiquitinated proteins and the proteasome are not linked to developmental defects, we also performed *akir-1* RNAi treatment starting from the L4 larval stage when cell division in the intestine has ceased (Altun & Hall, 2009). These intestinal polyubiquitin-binding reporter animals, examined at day 3 of adulthood, displayed the same distinctive converged fluorescent pattern (Figure 8D), as detected when RNAi treatment was started at L1 larval stage (Figure 1A).

**Figure 8.**
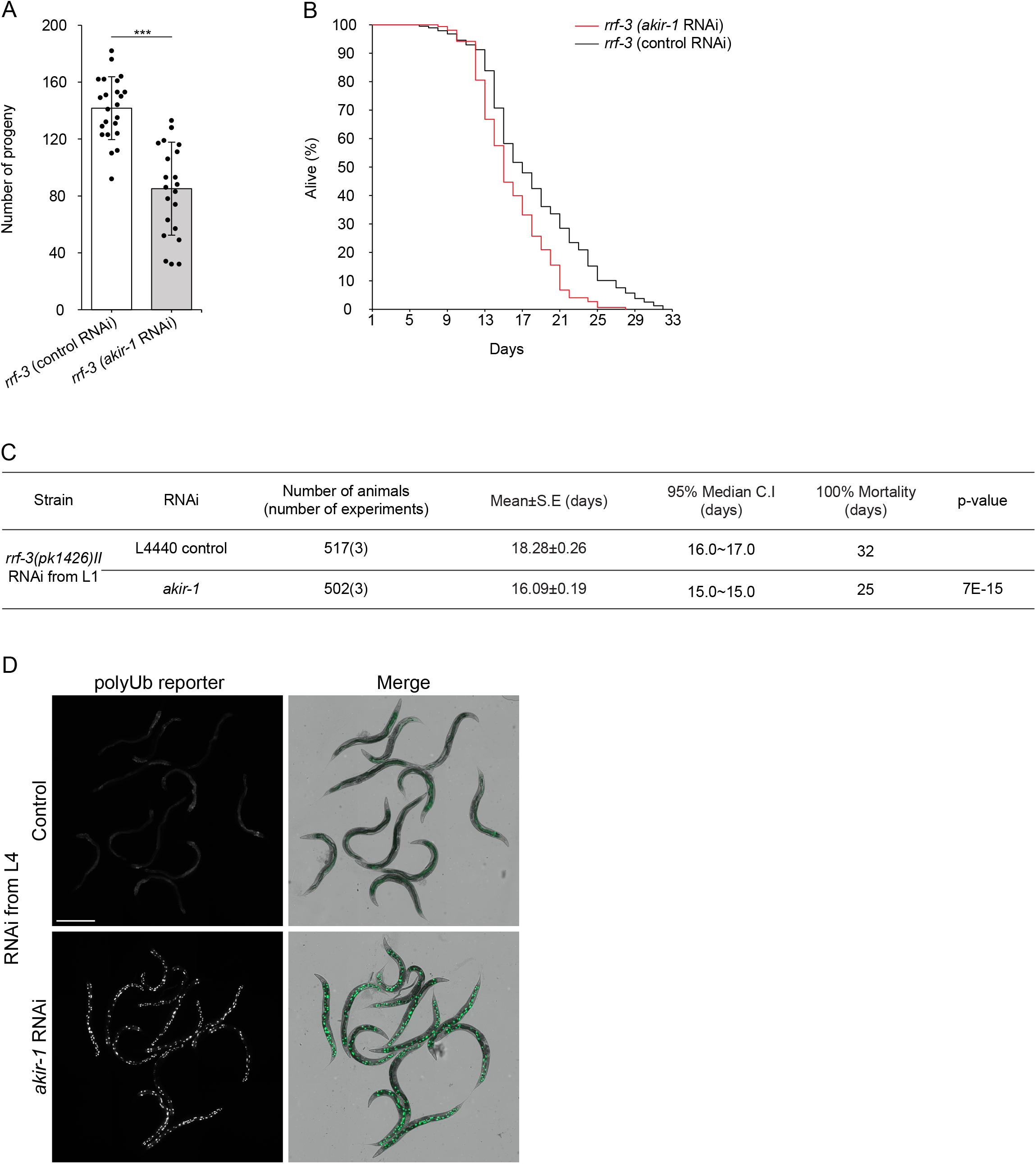
Downregulation of *akir-1* reduces progeny number and lifespan, and promotes nuclear accumulation of polyubiquitin-binding reporter even at adulthood. (**A**) The graph shows the mean number of progeny of *rrf-3(pk1426)* animals exposed to control or *akir-1* RNAi treatment, n = 3 independent experiments (total number of Po animals is 23 for control RNAi and 22 for *akir-1* RNAi). Error bars, SD; ***p < 0.001. (**B**) A representative Kaplan-Meier survival curve from one experiment with *rrf-3(pk1426)* animals exposed to control or *akir-1* RNAi treatment started at the L1 larval stage. (**C**) The table shows the statistics of three independent lifespan experiments of *rrf-3(pk1426)* animals treated with control or *akir-1* RNAi started at the L1 larval stage, and significance of *akir-1* RNAi treatment compared to control RNAi treatment determined with a Mantel-Cox (log-rank) test. Mean, restricted mean survival; C.l, confidence interval. (**D**) Representative fluorescence micrographs of 3-day adult animals expressing the polyubiquitin (polyUb) reporter in intestinal cells (*vha-6p::UIM2-ZsProSensor*) and treated with control or *akir-1* RNAi (left panels). Merge represents overlay of fluorescence and bright-field images (right panels). Scale bar, 500 µm.

We questioned by which molecular mechanisms AKIR-1 affects endogenous polyubiquitinated proteins and proteasomes. Previously, Akirins have been shown to act in the regulation of transcription via chromatin remodeling complexes (Bosch et al., 2020). In *C. elegans* AKIR-1 interacts specifically with the NuRD (NuRD I, II and MEC) chromatin remodeling complexes (Polanowska et al., 2018; Bowman et al., 2020), which have been reported to display some subunit overlap (Figure 8 ─ figure supplement 1A) (Passannante et al., 2010; Polanowska et al., 2018). These chromatin remodeling complex genes were not recognized as hits in our genome-wide RNAi screen, which could have been due to differences in experimental setup and fluorescence detection between the genome-wide scale and select gene RNAi, or due to the developmental criteria followed in our genome-wide RNAi screen. Thus, we next performed downregulation by select gene RNAi for each member of the NuRD complexes and MEC complex on the intestinal polyubiquitin-binding reporter animals, but detected no phenotypes mimicking the *akir-1* RNAi–induced effect (Figure ─ 8 figure supplement 1B). To directly affect transcription, we also performed RNAi against *ama-1*, a subunit of the RNA polymerase II, and while we observed a growth phenotype, the localization of polyubiquitin-binding reporter was not affected (Figure 8 ─ figure supplement 1C).

As our results show an accumulation of polyubiquitinated proteins in conjunction with a reduction in proteasome levels in the intestinal nuclei upon AKIR-1 depletion, one explanation could be an involvement of AKIR-1 in the nuclear import of proteasomes. The nuclear transport machinery is required for protein trafficking between the cytoplasm and the nucleus, and members of the karyopherin family function as cargo receptors. In *C. elegans*, 13 members of the karyopherin family have been identified (Figure 9A) (Adam, 2009). We performed RNAi against all these karyopherin family members using our intestinal polyubiquitin-binding reporter strain (Figure 9B, Figure 9 – figure supplement 1). Of these, downregulation of importin α3, *ima-3*, showed clearly a similar polyubiquitin phenotype as *akir-1* RNAi (Figure 9B), although the intestinal fluorescent puncta formed with weaker intensity. DNA staining with Hoechst confirmed that the polyubiquitin-binding reporter concentrated to the intestinal nuclei upon *ima-3* RNAi treatment (Figure 9C). In addition, downregulation of importin β1, *imb-1*, partially mimicked the *akir-1* RNAi–induced fluorescence phenotype, but to a lesser extent than *ima-3* RNAi. (Figure 9 ─ figure supplement 1B), and with an apparent sick phenotype. We next tested for genetic interactions between *akir-1* and *ima-3* by performing *ima-3* RNAi on *akir-1* mutants (Figure 9D). Compared to either *akir-1* mutants or *ima-3* RNAi–treated wild-type animals, the *akir-1* mutant animals exposed to *ima-3* RNAi showed severely delayed growth and reduced body size (Figure 9D), suggesting that AKIR-1 and IMA-3 may cooperate in some cellular processes. Whilst it has been suggested that individual importins act as receptors for several different types of protein cargo (Kimura et al., 2017), we reasoned that AKIR-1 is likely involved in a more limited role. We therefore used a nuclear localized GFP transgenic reporter strain (*sur-5p::NLS-GFP*) (Yochem et al., 1998; Mikkonen et al., 2017) to investigate whether AKIR-1 affects nuclear import in general. As expected, RNAi against *ima-3* markedly reduced the GFP signal in intestinal nuclei, but *akir-1* RNAi had no effect on the nuclear localization of the GFP reporter suggesting a more specific role for *akir-1* in nuclear import than for importins (Figure 9E). Taken together, our genetic experiments suggest that AKIR-1 acts in the nuclear import of proteasomes cooperating with importins α3 and β1.

**Figure 9.**
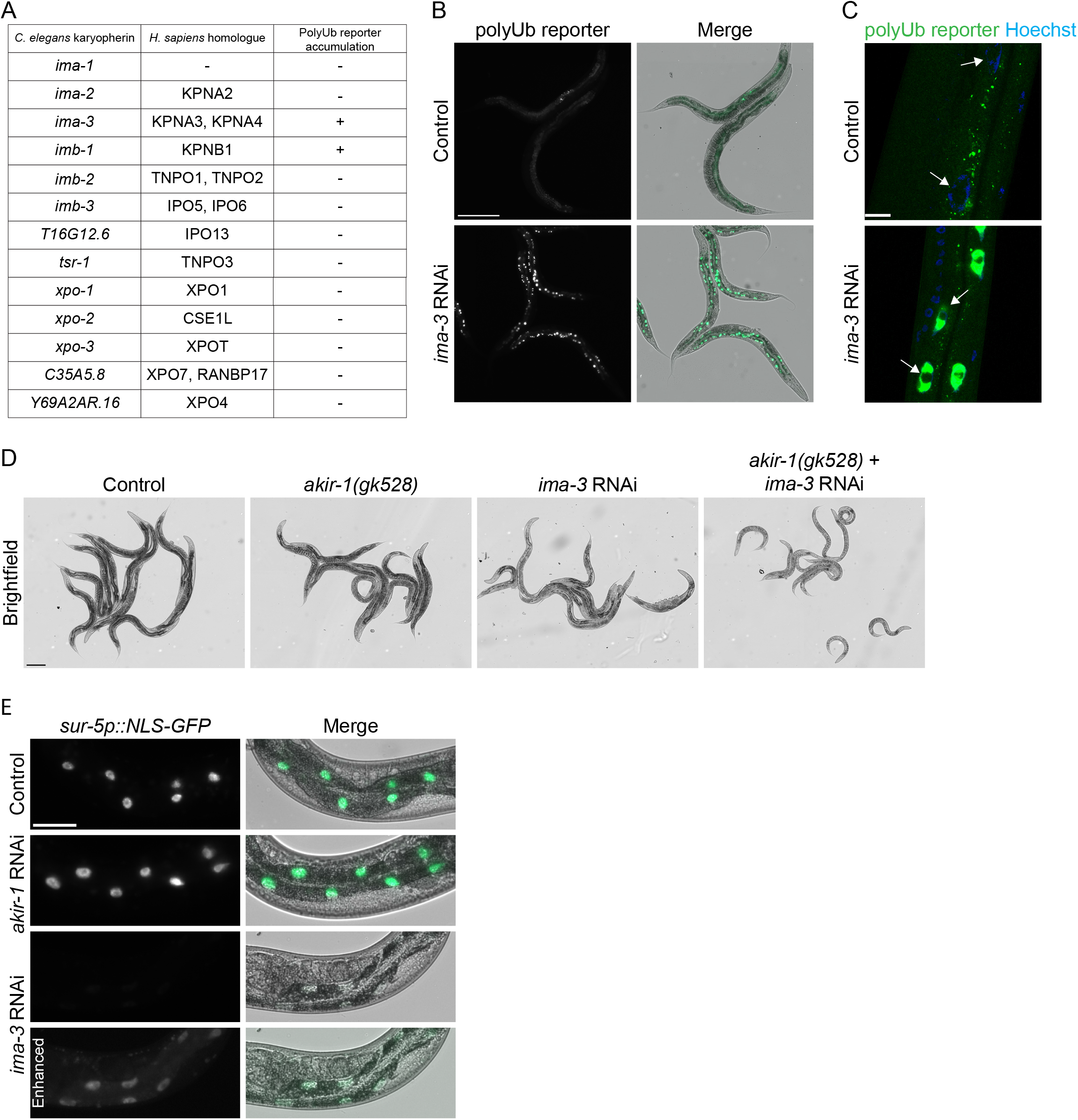
Loss of the *akir-7*-genetic interactor *ima-3* induces nuclear accumulation of polyubiquitinated proteins, and *akir-1* is not required for general nuclear import. (**A**) The table shows *C. elegans* karyopherins, their human orthologues, and RNAi-induced nuclear accumulation of the polyubiquitin (polyUb) reporter. (**B**) Representative fluorescence micrographs of control and *ima-3* RNAi-treated N2 animals expressing the polyubiquitin (polyUb) reporter (*vha-6p::UIM2-ZsProSensor*) in the intestinal cells (left panels). Merge represents overlay of fluorescence and bright-field images (right panels). Scale bar, 200 µm. (**C**) Representative confocal micrographs of control and *ima-3* RNAi-treated polyubiquitin reporter animals with Hoechst-visualized nuclei. White arrows point to representative intestinal cell nuclei. Scale bar, 10 µm. (**D**) Representative bright-field images of control (N2) and *akir-1 (gk528)* mutant animals treated with control or *ima-3* RNAi, respectively. Scale bar, 200 µm. (**E**) Representative fluorescence micrographs of control, *akir-1*, and *ima-3* RNAi-treated wild-type animals expressing ubiquitiously nuclear-localized GFP (*sur-5p::NLS-GFP*) (left panels). Merge represents overlay of fluorescence and bright-field images (right panels). Lowest panels show enhanced fluorescence signal. Scale bar, 50 µm.

## Discussion

Our results are the first to show the importance of AKIR-1 for nuclear localization of the proteasome and protein homeostasis in a multicellular organism. The *C. elegans* AKIR-1 belongs to the highly conserved, metazoan-specific Akirin family of small nuclear proteins. The number of Akirin paralogues within species varies from one to eight, with *C. elegans* having one copy, and vertebrates having two, AKIRIN1 and AKIRIN2 (Artigas-Jeronimo et al., 2018; Bosch et al., 2020). Akirins are best known for their roles in innate immunity, where they are required for expression of downstream effector genes in several organisms (Artigas-Jeronimo et al., 2018). In addition, Akirins are involved in developmental processes (Bosch et al., 2020).

Recently, a screen for regulators of the levels of the transcription factor MYC by de Almeida and colleagues identified an additional function for human AKIRIN2 as a regulator of the turnover of a subset of nuclear proteins in human cancer cells (de Almeida et al., 2021). As downregulation of *AKIRIN2* led to reduced nuclear fluorescence signal of tagged proteasome subunits, and as AKIRIN2 was shown with cryo-EM to bind to the 20S proteasome, they suggested that AKIRIN2 acts as a mediator of proteasome nuclear import. We identified the *AKIRIN2* orthologue, *akir-1*, in a genome-wide RNAi screen for novel proteasome regulators in *C. elegans*. Here, we show that AKIR-1 is required for nuclear localization of endogenous proteasomes. More precisely, we demonstrate that animals lacking *akir-1* have reduced levels of nuclear proteasomes, both in intestinal cells and oocytes. Thus, our results reveal the significance of Akirins in regulating nuclear proteasome localization at an organismal level, and that this function is conserved between human cells and invertebrates.

The molecular mechanism behind Akirins’ many physiological functions is believed to be transcriptional regulation due to their interaction with chromatin remodeling complexes and transcription factors (Goto et al., 2008; Nowak et al., 2012), and as such the effect of AKIR-1 on proteasome localization could be indirect. However, we observed no phenotype resembling the *akir-1* RNAi–induced phenotype upon downregulation of all subunits of the *C. elegans* NuRD and MEC chromatin remodeling complexes, as well as a subunit of RNA polymerase II. A transcription-independent mechanism of AKIR-1 is further supported by our Western blot analysis showing no consistent change in proteasome levels upon *akir-1* depletion, and by a report describing a post-transcriptional effect on the meiosis-linked SYP-1 protein in the *akir-1* mutants (Clemons et al., 2013). Thus, we reasoned that AKIR-1 could directly control nuclear transport of proteasomes in *C. elegans*.

Nuclear transport of proteins is mediated by the importins and exportins of the karyopherin family. We show that knockdown of two importin family members, *ima-3* and *imb-1* mimic the *akir-1* RNAi– induced nuclear accumulation of the polyubiquitin-binding reporter in intestinal cells. The small fluctuation in observed phenotypes between *akir-1* and *importins* knockdowns might derive from variations in protein stability, receptor redundancy, or the existence of more complex proteasome transport mechanisms. We further show that *akir-1* genetically interacts with *ima-3*. Previously, it has been reported that GFP-tagged AKIR-1 interacts with IMA-3 (Polanowska et al., 2018), and that AKIR-1 binds to IMA-2 in a yeast two-hybrid screen (Bowman et al., 2019). Interestingly, the *S. cerevisiae* importin α family member SRP1, which is believed to be a homologue of *C. elegans ima-2* and *ima-3*, is required for nuclear import of proteasomes (Lehmann et al., 2002; Adam, 2009). Importin 9 (IPO9) has also been implicated to play a similar role in *D. melanogaster* during spermatogenesis and in human cancer cells (de Almeida et al., 2021; Palacios et al., 2021), but *C. elegans* has no IPO9 homologue (Adam, 2009). Our results suggest that AKIR-1 controls proteasome nuclear import in *C. elegans* together with importin receptors IMA-3 and IMB-1.

In addition to direct interaction between importins and proteasome subunits, it has been shown in yeast that adaptor proteins, such as Sts1 in *S. cerevisiae* and its orthologue Cut8 in *S. pombe*, interact with the 20S/26S proteasome (Tatebe & Yanagida, 2000; Takeda & Yanagida, 2005; Chen et al., 2011; Takeda et al., 2011; Budenholzer et al., 2020). A homologue of Sts1/Cut8 has so far been found only in *D. melanogaster* (Takeda & Yanagida, 2005), but de Almeida and colleagues hypothesized that the human AKIRIN2 could potentially be a functional homologue of Sts1/Cut8, even though lack of sequence homology (de Almeida et al., 2021). Both AKIRIN2 and Sts1 contain protein regions of predicted disorder and are short-lived proteins (Budenholzer et al., 2020; de Almeida et al., 2021). Our *akir-1* RNAi experiments, starting at L1 or L4 larval stage, consistently displayed a highly penetrant nuclear phenotype in the intestinal polyubiquitin-binding reporter strain, suggesting that AKIR-1 is a short-lived protein also in *C. elegans* intestine. Our results provide further support that Akirins might function as adaptor proteins in the nuclear transport of proteasomes.

Our study further demonstrates a broader role of AKIR-1 in regulation of proteasome function and protein homeostasis in a multicellular organism. Lysates of *akir-1* RNAi–treated animals contained slightly increased *in vitro* proteasome activity, which was mainly due to enhanced activity of the 20S particle. Interestingly, no change was detected in proteasomal degradation of the photoconverted UbG76V-Dendra2 reporter at the cellular level in living animals in either intestinal cells, or body-wall muscle cells, suggesting that the increase in *in vitro* proteasome activity does not stem from a general systemic response to *akir-1* knockdown. Remarkably, when we determined proteasomal degradation rate at the subcellular level, *i*.*e*., by separately measuring degradation in the nucleus and cytoplasm of intestinal cells, the *akir-1* RNAi animals showed a clear increase in the nuclear to cytoplasmic fluorescence ratio of photoconverted UbG76V-Dendra2, revealing that downregulation of *akir-1* slows proteasomal degradation in the nucleus. This reduced degradation capacity in the nucleus reflects the decreased nuclear localization of the proteasome.

We have previously reported tissue-specific variations in proteasome activity and regulatory mechanisms in *C. elegans* (Hamer et al., 2010; Matilainen et al., 2013; Pispa et al., 2020). Interestingly, here we show that although oocytes display a more pronounced reduction in the nuclear proteasome levels compared to intestinal cells, acute accumulation of endogenous polyubiquitinated proteins is not induced in the nuclei of these cells upon loss of *akir-1*. Cell type– specific differences in proteasomal substrates could potentially contribute to the different response in the nuclear accumulation of polyubiquitinated proteins between oocytes and intestinal cells. It has also been reported that dissected gonads from young *D. melanogaster* flies display elevated proteasome capacity compared to somatic tissues (Fredriksson et al., 2012; Tsakiri et al., 2013), which could contribute to a faster protein turnover in oocytes. Additionally, it has been reported that ubiquitinated proteins are exported from the nucleus to the cytoplasm in proteasome inhibitor– treated cell lines (Hirayama et al., 2018). In our study, the *akir-1* depletion–induced reduction in nuclear proteasomes could mimic proteasome inhibition at this subcellular compartment, and thereby potentially result in nuclear export of polyubiquitinated proteins in oocytes. Accordingly, we occasionally observed an accumulation of polyubiquitin-positive staining at the rim of the nuclear membrane in oocytes of *akir-1* depleted animals. The underlying mechanisms of this finding should be explored in future studies.

In addition to oocytes, body-wall muscle cells also responded differently to *akir-1* knockdown compared to the intestinal cells. In the body-wall muscle cells, we observed a later onset of the phenotype, as a slight nuclear accumulation of polyubiquitin-binding reporter was detected only in the next (F_1_) generation after continuous exposure to *akir-1* RNAi treatment. It is unlikely that this later onset is caused by a variation in RNAi efficiency based on the use of a RNAi-sensitive strain and as we have previously shown efficient RNAi capacity in both body-wall muscle cells and intestinal cells (Mikkonen et al., 2017). This could be due to tissue-specific variation in proteasomal degradation rate, as we have demonstrated slower degradation in the body-wall muscle cells compared to intestinal cells (Hamer et al., 2010; Matilainen et al., 2013). We speculate that due to the slower proteasomal degradation rate, body-wall muscle cells maintain functional AKIR-1 proteins longer after *akir-1* knockdown compared to intestinal cells. Given the variable effects of AKIR-1 depletion, future studies are required to decipher the contribution of individual molecular mechanisms to the tissue-specific phenotypes. Overall, our AKIR-1 study demonstrates that the role of Akirins in regulating nuclear proteasome localization is conserved between *C. elegans* and human cells, and that Akirin family members can interact with several nuclear transport proteins. Lastly, and most importantly, our study suggests a broader role for Akirins in healthspan regulation and maintenance of cellular protein homeostasis in a tissue-specific manner in multicellular organisms.

## Materials and Methods

### C. elegans strains

All strains were cultured as previously described (Brenner, 1974) at 20 °C on nematode growth medium (NGM) plates seeded with OP50 *Escherichia coli* bacteria. N2 Bristol, VC1056[*akir-1(gk528)I*], and NL2099[*rrf-3(pk1426)II*] strains were obtained from the Caenorhabditis Genetic Center (CGC, Minneapolis, MN, USA). The *akir-1* mutant allele *gk528* was backcrossed three times with wild-type N2 animals. Following strains were also used: YD3[*xzEx3[unc-54p::UbG76V::Dendra2]*] (Hamer et al., 2010), YD27[*xzEx27[vha-6p::UbG76V::Dendra2]*] (Li et al., 2011), YD90[*xzIs1[vha-6p::UIM2::ZsProSensor]*] (Matilainen et al., 2013), GR1702[*Is1[sur-5p::NLS-GFP]*] (Mikkonen et al., 2017), CER395*[ubh-4(cer68[ubh-4::egfp])II]* (Vicencio et al., 2019), YD116[*rrf-3(pk1426);xzIs2[unc-54p::UIM2::ZsProSensor]*] (Jha & Holmberg, 2020), and YD114[*xzIs2[unc-54p::UIM2::ZsProSensor]*] (Pispa et al., 2020). To generate YD117[*xzEx113[rpt-5p::GFP::RPT-5]*] reporter strain, GFP::RPT-5 translational fusion construct was created by substituting the let-858 promoter of the GFP vector pPD118.25 (L3786 was a gift from Andrew Fire; Addgene plasmid #1593; http://n2t.net/addgene:1593; RRID:Addgene_1593; Watertown, MA, USA) with a 2010-bp putative rpt-5 promoter sequence, mutating the GFP stop codon, and replacing the let858 3’ UTR with a 1747-bp sequence covering the 1504-bp rpt-5 coding region with introns and the following 183-bp 3’ UTR region. To create extrachromosomal transgenic lines, the plasmid DNA (120-180 ng/µl) was injected into gonads of young N2 adults along with *P*_*myo-2*_ ::CFP (20 ng/ µl) marker. Unless otherwise stated, animals were age-synchronized with bleach treatment, and L1 larvae (day 1) were placed on bacteria-seeded plates prior being collected at first day of adulthood (day 4). Due to slow growth age-synchronized *akir-1* mutant L1 larvae were placed on bacteria-seeded plates 10-17 hours before the age-synchronized N2 L1 larvae. Both N2 and *akir-1* mutant animals were collected at first day of adulthood (day 4 with N2 and day 5 with *akir-1* mutant animals). The experiments were done either with a hermaphrodite population, with males existing at a frequency of <0.2% in N2 (Hodgkin & Doniach, 1997) and 0,9% in *akir-1(gk528)* (Clemons et al., 2013), or with singly picked hermaphrodites.

### C. elegans RNA interference (RNAi)

Select gene RNAi was performed using the feeding method as described earlier (Timmons et al., 2001) with a few changes. The HT115(DE3) bacterial strain carrying the empty *pL4440* cloning vector was used as a control. Bacterial strains were cultured overnight at 37 °C in Luria broth (LB) medium containing 100 μg/ml ampicillin (Merck KGaA Cat#A0166, Darmstadt, Germany) and 12.5 μg/ml tetracycline (Merck KGaA Ca#T7660). The culture was diluted 100-fold and allowed to grow in 2xYT medium containing ampicillin and tetracycline in above-mentioned concentrations until the absorbance at 600 nm (OD600) reached 0.4. The double-stranded RNA expression was induced using 0.4 mM isopropyl β-D-1-thiogalactopyranoside, IPTG (Merck KGaA Ca#I6758). Before adding bacterial cells on RNAi feeding agar plates, cultures were supplemented with additional ampicillin and tetracycline, and the IPTG concentration was increased to 0.8 mM. RNAi feeding agar plates were composed of standard NGM complemented with 100 μg/ml ampicillin, 12.5 μg/ml tetracycline, and 0.4 mM IPTG. For experiments related to quantification of changes in fluorescence of YD90, YD114 or YD116 animals, the RNAi clones were cultured for six hours at 37 °C in 2xYT medium complemented with the above-mentioned antibiotics, induced with 0.8 mM IPTG, and cultured overnight at 37 °C prior to plating on the RNAi feeding agar plates. Unless otherwise stated, animals were age-synchronized with bleach treatment, and L1 larvae (day 1) were placed on control or RNAi-seeded feeding plates prior being collected at first day of adulthood (day 4). When *akir-1* mutant animals were exposed to RNAi treatment, age-synchronized L1 larvae were placed on the RNAi plates 10-17 hours before the age-synchronized N2 L1 larvae. Both N2 and *akir-1* mutant animals were collected at first day of adulthood (day 4 with N2 and day 5 with *akir-1* mutant animals). RNAi clones for the following genes were used: *akir-1(E01A2*.*6), hda-1(C53A5*.*3), let-418(F26F12*.*7), lin-53(K07A1*.*12), lin-40(T27C4*.*4), chd-3(T14G8*.*1), mep-1(M04B2*.*1), dcp-66(C26C6*.*5), ama-1(F36A4*.*7), ima-1(T19B10*.*7), ima-2(F26B1*.*3), ima-3(F32E10*.*4), imb-2(R06A4*.*4), imb-3(C53D5*.*a), T16G12*.*6, tsr-1(F53G2*.*6), xpo-1(ZK742*.*1), xpo-2(Y48G1A*.*5), xpo-3(C49H3*.*10)* (J. Ahringer RNAi library, Source BioScience, Nottingham, UK), or *imb-1(F28B3*.*8), C35A5*.*8, Y69A2AR*.*16* (clones were kind gifts from Dr. Susana Garcia; Vidal ORFeome). All phenotype-inducing RNAi clones as well as most of the no phenotype–inducing clones were confirmed by sequencing (Eurofins Genomics, Ebersberg, Germany).

The experimental setup of our original genome-wide RNAi screen for UPS regulators was slightly different than for the select gene RNAi. Briefly, the J. Ahringer RNAi library clones were cultured overnight at 37 °C in 2xYT medium containing antibiotics without a subsequent culture with IPTG before seeding. Age-synchronized YD90[*xzIs1[vha-6p::UIM2::ZsProSensor]*] (Matilainen et al., 2013) L1 larvae were placed on RNAi seeding plates, and manually scored after three days as young adults for visible changes in fluorescence with Leica MZ16 FA fluorescence stereomicroscope with a GFP Plus filter (Leica Microsystems, Wetzlar, Germany). Animals with delayed or arrested development or otherwise atypical appearance were censored from further study. Further, RNAi clones affecting general protein synthesis, monitored by changes in the fluorescence of YD25*[xzEx25[vha-6p::Dendra2]]* strain (Li et al 2011), were excluded as hits.

### Quantitative real-time PCR (qPCR)

Age-synchronized animals treated with RNAi were harvested in M9 buffer (22 mM KH_2_ PO_4_, 41 mM Na_2_ HPO_4_, 8,5 mM NaCl, and 19 mM NH_4_ Cl) at first day of adulthood and stored at –80 °C. RNA was extracted using Nucleospin RNA kit (MACHEREY-NAGEL GmbH & Co. KG Cat#740955, Düren, Germany) and cDNA synthesis was done with Maxima First Strand cDNA synthesis kit for RT-qPCR (Thermo Fisher Scientific Cat#K1641, Waltham, MA, USA). Quantitative real-time PCR was performed with Maxima SYBR Green/Rox qPCR Master Mix (Thermo Fisher Scientific Cat# K0221) and LightCycler 480 quantitative PCR machine (Roche Diagnostics International AG, Rotkreuz, Switzerland). The qPCR data were normalized to the expression of three reference genes (*act-1, cdc-42*, and *pmp-3*), and comparative Ct (ΔΔCt) method was used to quantify relative expressions of *akir-1* mRNA. Oligo sequences were as follows: 5’-gatatgcgaacgtctgctca, 5’-ggaatagtcatccccagtgc (*akir-1*); 5’-tcggtatgggacagaaggac, 5′-catcccagttggtgacgata (*act-1*); 5’-ctgctggacaggaagattacg, 5′ -ctcggacattctcgaatgaag (*cdc-42*); and 5′-gttcccgtgttcatcactcat, 5′-acaccgtcgagaagctgtaga (*pmp-3*).

### Gel electrophoresis followed by Western immunoblot analysis or in-gel proteasome activity assay

Age-synchronized animals fed with OP50 or RNAi bacteria were harvested in M9 buffer at first day of adulthood and animal pellets were stored at -80 °C. Pellets for immunoblot analysis were lysed using lysis buffer (50 mM Hepes, 150 mM NaCl, 5 mM EDTA) supplemented with 20 mM N-ethylmaleimide, NEM (Merck KGaA Cat# E3876) and 10 μM MG-132 (Peptides International Cat# IZL-3175-v, Louisville, KY, USA) to inhibit deubiquitination and degradation of ubiquitin-conjugated proteins, respectively. In addition, lysis buffer was supplemented with a Pierce Protease Inhibitor Mini Tablet (Thermo Fisher Scientific Cat# A32955). Lysed samples were separated on a SDS (sodium dodecyl sulfate) polyacrylamide gel and blotted onto a nitrocellulose membrane using Trans-Blot Turbo Transfer System (Bio-Rad Laboratories, Hercules, California, USA). Antibodies against proteasome 20S α-subunits, MCP231 (Enzo Life Sciences Cat# BML-PW8195, RRID:AB_10541045, New York, NY, USA) in 1:1000 dilution, polyubiquitinated proteins, FK1 (StressMarq Biosciences Cat# SMC-213, RRID:AB_2699340, Victoria, Canada) in 1:500 dilution, and alpha-tubulin (Sigma-Aldrich Cat# T5168, RRID:AB_477579) in 1:10000 dilution were used in blotting. The following HRP-conjugated anti-mouse IgM (Millipore Cat# 401225-2ML, RRID:AB_437770) and IgG (Promega Cat# W4021, RRID:AB_430834, Madison, WI, USA) antibodies in 1:10 000 dilutions were used. ECL signals were visualized with Odyssey FC Imaging System (LI-COR, Lincoln, NE, USA), and quantified with Image Studio software (Li-COR). Alpha-tubulin signal was used for normalization.

Native gel electrophoresis and the in-gel proteasome activity assay were performed as earlier reported with a few exceptions (Elsasser et al., 2005). Animal pellets were lysed using a dounce homogenizer and native gel lysis buffer (Elsasser et al., 2005). Gels were run in an ice bath for 30 minutes at 20 mA and then for 2 hours at 40 mA. The gels were incubated in developing buffer containing 160 μM of fluorogenic proteasome substrate succinyl-leu-leu-val-tyr-7-amino-4-methylcoumarin, suc-LLVY AMC (Bachem Cat# l.1395, Bubendorf, Switzerland) and imaged either with MultiImage Light Cabinet using FluorChem 8900 software (Alpha Innotech Corporation, San Leandro, CA, USA) or Gel Doc XR+ System with Image Lab Software (Bio-Rad). Coomassie staining with Colloidal Blue staining kit (Thermo Fisher Scientific Cat# LC6025) was used for normalization and image analysis were made with Fiji software (Schindelin et al., 2012).

### C. elegans immunofluorescence with dissected animals

Age-synchronized animals cultured either on OP50 seeded plates or RNAi feeding plates were harvested at first day of adulthood in M9 buffer. Animals were transferred onto a glass dish and dissected using 27-gauge syringe needles. To immobilize the animals 1 mM levamisole hydrocloride (Merck KGaA Cat# T7660) was used prior making incision close to the pharynx forcing the intestine and gonad to extrude. Dissected animals were fixed with 2xRFB (160 mM KCL, 40 mM NaCl, 20 mM EGTA, 10 mM spermidine, 30 mM PIPES pH 7.4, 50% methanol, and 1% formaldehyde). Additional 100% methanol fixation for 1 min was utilized when antibody against the proteasome 20S alpha subunits was used. Fixed dissected animals were permeabilized using 0.5 or 1% Triton X-100 in PBS and mounted with SlowFade Diamond Antifade Mountant (Thermo Fisher Scientific Cat# S36967). Antibodies against polyubiquitinated proteins, FK1 (RRID:AB_2699340) and the proteasome 20S alpha subunits, MCP231 (RRID:AB_10541045) were used in 1:200 dilution. The following Alexa fluor 594 conjugated anti-mouse IgM (Thermo Fisher Scientific Cat# A-21044, RRID:AB_2535713) and IgG (Thermo Fisher Scientific Cat# A-11005, RRID:AB_2534073 or Cat# R37121, RRID:AB_2556549) secondary antibodies in 1:200 and 1:100 dilution respectively were used for visualization. The nuclear immunostaining was visually estimated as none, when nuclear staining intensity was similar or less compared to staining intensity in the cytoplasm. A higher nuclear staining intensity compared to the cytoplasmic staining was further estimated visually as weak or strong. DNA was stained with 4 ug/ml Hoechst 33342 (Merck KGaA Cat# B2261).

### Lifespan Assays and Progeny Counts

Lifespan experiments were performed at 20 °C. Age-synchronized animals were plated on RNAi feeding plates as L1 larvae (day 1). Animals were transferred to a new plate, first every second day, and then every few days after they stopped producing offspring. Animals were checked daily, and animals failing to respond to a gentle prod with a platinum worm pick were classified as dead. Animals crawling off the plate, dying of an extruded gonad, or carrying internally hatched offspring were censored at the time of their death. Basic survival analysis was performed using an online tool, OASIS 2 (Han et al., 2016). Data from all three separate lifespan experiments were combined and used for analysis as one data set. For counting progeny, age-synchronized animals were plated individually on RNAi feeding plates as L1 larvae. Animals were moved to fresh RNAi feeding plates every day, until they stopped producing offspring. Total viable offspring per animal was counted, censoring the offspring of animals that crawled off the plate, died of internally hatched offspring or an extruded gonad before finishing egg-laying.

### Microscopy

Age-synchronized animals were imaged at first day of adulthood unless otherwise mentioned. Animals were mounted on 3% agarose pads and immobilized with 1 mM levamisole. Group of 5 to 15 live animals were imaged with a Zeiss Axio Imager 2 upright wide-field light microscope and a Zeiss EC Plan Neofluar numerical aperture (NA) 10 × 0.3 objective or a Plan Apochromat NA 20 × 0.8 objective, or with a LSM 780 inverted confocal microscope and a Plan-Neofluor NA 40 × 1.3 or a Plan-Apochromat NA 63 × 1.40 objective, or with a Zeiss LSM 880 inverted confocal microscope and a Plan-Apochromat NA 40 × 1.40 or NA 63 × 1.4 objective (Zeiss, Oberkochen, Germany). All the microscopes run a Zeiss Zen 2 software (Zeiss). For photoconversion of the tissue-specific UbG76V-Dendra2, green Dendra2 protein was converted to red using 405-nm UV light. Degradation of the red signal was followed 16-18 (intestine) or 24 (muscle) hours later. For determining the nuclear/cytoplasmic ratio of the red UbG76V-Dendra2 signal, images were taken with the Zeiss LSM 880 inverted confocal microscope and the Plan-Apochromat NA 63 × 1.4 objective, and mean fluorescent intensity of both the nuclei and cytoplasm were quantified using Microscopy Image Browser software (Belevich et al., 2016).

Fluorescence signal was quantified from the original, unmodified TIFF file format images with Fiji software. In Fiji, sliding paraboloid algorithm was used to subtract background. The mean fluorescence intensity was quantified using an adequate threshold to select fluorescence signal and using the measure function. The brightness of the fluorescence signal has been adjusted in some images using Fiji or Adobe Photoshop (Adobe, San Jose, CA, USA), and all images presented together for comparison were adjusted similarly. For Hoechst nuclear staining, age-synchronized animals were washed with M9, fixed for 2 min using 100% methanol, and permeabilized either with 0,5% or 1% Triton-X-100 in PBS. DNA was labeled with 4 ug/ml Hoechst 33342 stain (Merck KGaA Cat# B2261).

### Statistical analysis

The statistical significance was determined using Welch’s t-test (two-tailed distribution and unequal variance) using Microsoft Excel 2016 spreadsheet (Microsoft, Redmond, WA, USA). For lifespan analysis, statistical significance was determined with a Mantel-Cox (log-rank) test using RStudio software (RStudio, Boston, MA, USA) and the lifespan data used was stratified into three independent experiments.

## Supporting information

Supplementary figures v2 Pispa Mikkonen et al 2022

## Acknowledgements

This study was supported by grants to C.I.H from the Academy of Finland (259797, 297776), Sigrid Jusélius Foundation, Medicinska Understödsföreningen Liv och Hälsa r.f., and Magnus Ehrnrooth Foundation. E.M. was supported by the Doctoral Programme in Biomedicine of University of Helsinki and by grants from the Finnish Cultural Foundation. J.C. lab was funded by Ministerio de Ciencia, Innovación y Universidades, which is part of Agencia Estatal de Investigación (AEI), through the Retos grant (PID2020-114986RB-100). The authors thank the members of the Biomedicum Imaging Unit (BIU) core facility of University of Helsinki for help with imaging, Dr. Ilya Belevich from the Electron Microscopy Unit (EMBI) core facility, University of Helsinki, for help with the Microscopy Image Browser software, Dr. Susana Garcia for sharing a *C. elegans* strain and RNAi clones, and Holmberg lab members Anni-Helena Sukupolvi and Sweta Jha for assistance with initial experiments. Some strains were provided by the CGC, which is funded by NIH Office of Research Infrastructure Programs (P40 OD010440). VC1056 strain was provided by the *C. elegans* Reverse Genetics Core Facility at the University of British Columbia, which is part of the international *C. elegans* Gene Knockout Consortium. The authors acknowledge networking support by the COST Action 1BM408/GENiE.

