## Supplementary figures v2 Pispa Mikkonen et al 2022 for "AKIR-1 Regulates Proteasome Localization and Function in *Caenorhabditis elegans*"

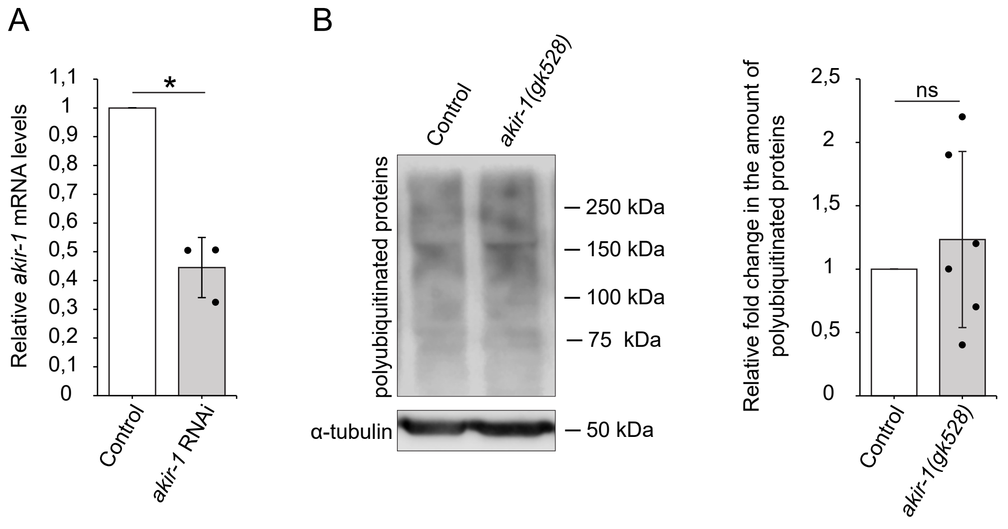


**Figure 2 — figure supplement 1.** RNAi treatment decreases *akir-1* mRNA levels and the total amount of polyubiquitinated proteins is not consistently changed in *akir-1* deletion mutants. (**A**) The graph shows average fold change in *akir-1* mRNA levels in *akir-1* RNAi–treated wild-type (N2) animals compared to control RNAi–treated animals (set as 1). n = 3 independent experiments; error bars, SD; *p < 0,05. (**B**) Western blot of polyubiquitinated proteins (left) in whole-animal lysates of control (N2) and *akir-1(gk528)* deletion animals. Anti-alpha-tubulin antibody was used as a normalization control. The quantification graph (right) shows the average fold change in *akir-1(gk528)* mutants compared to control (N2) animals (set as 1). n = 6 independent experiments; error bars, SD; ns, not significant.

A

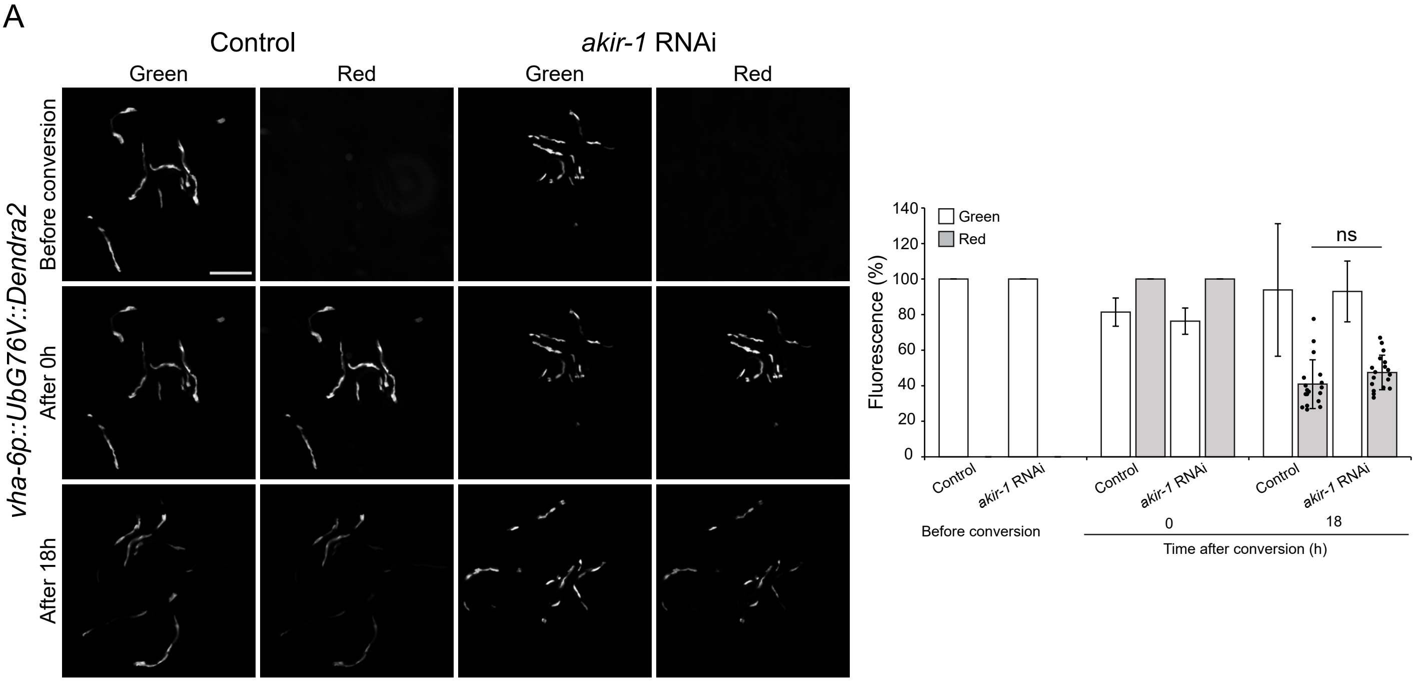


**Figure 4 — figure supplement 1.** Downregulation of *akir-1* does not change the degradation of UPS reporter proteins in the intestinal cells. (**A**) Representative fluorescence micrographs of green and red fluorescence of control and *akir-1* RNAi–treated transgenic animals expressing photoconvertible UbG76V-Dendra2 (*vha-6p::UbG76V::Dendra2)* reporter proteins in the intestinal cells before or 0 h and 18 h after whole-worm photoconversion. Scale bar, 500 µm. Note, this is an extended figure for Figure 4A including the red fluorescence images before conversion as well as all corresponding green fluorescent images. The graph shows the mean percentages of green and red fluorescence relative to the initial intensity (Before conversion) or at the indicated time points after conversion (0 h and 18 h). n = 6 independent experiments with triplicate images of 6-7 animals per image (total number of animals is 108); Error bars, SD; ns, not significant.

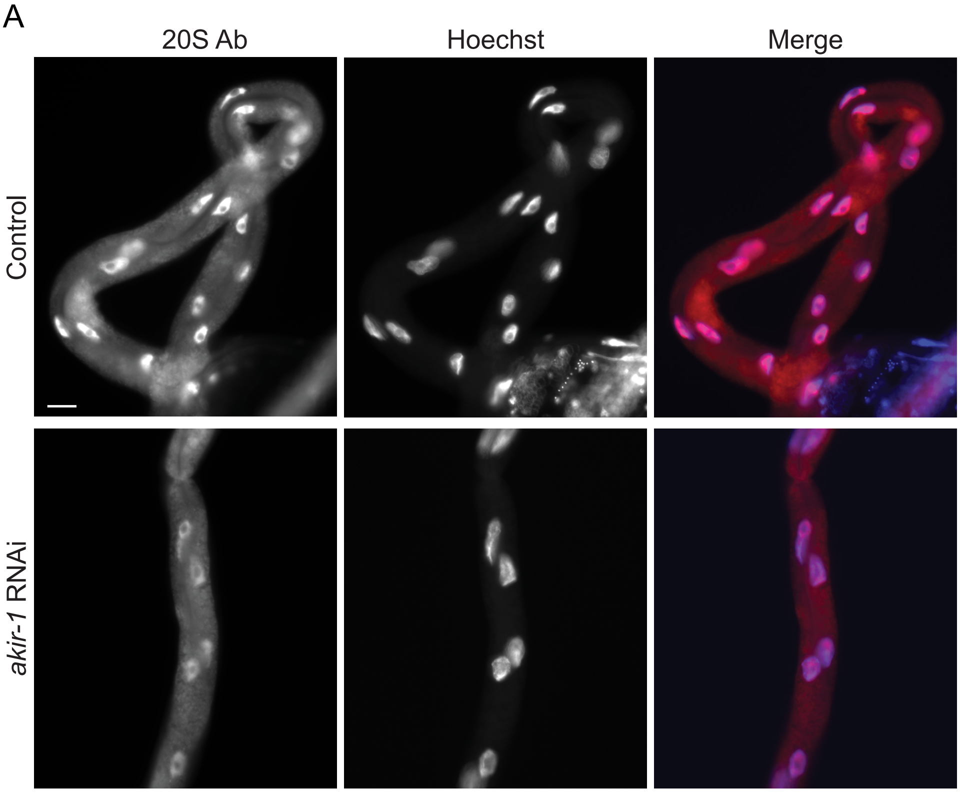


**Figure 5 — figure supplement 1.** Proteasome expression decreases in the nuclei of intestinal cells upon *akir-1* downregulation. (**A**) Representative micrographs of proteasome immunostaining (20S Ab) in dissected intestines of control and *akir-1* RNAi–treated wild-type animals. Nuclei visualized with Hoechst. Scale bar, 20 µm.

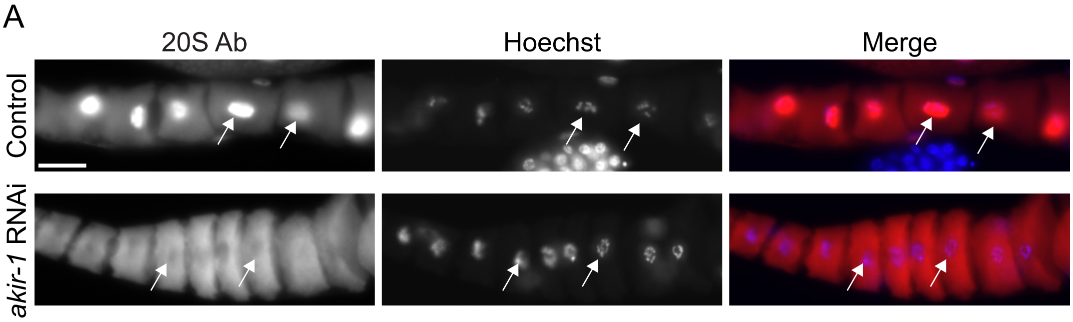


**Figure 6 — figure supplement 1.** Nuclear proteasome expression decreases in oocytes upon *akir-1* downregulation. (**A**) Representative micrographs of proteasome immunostaining (20S Ab) in dissected oocytes of control and *akir-1* RNAi–treated wild-type animals. Nuclei are visualized with Hoechst. White arrows point to representative nuclei. Scale bar, 20 µm.

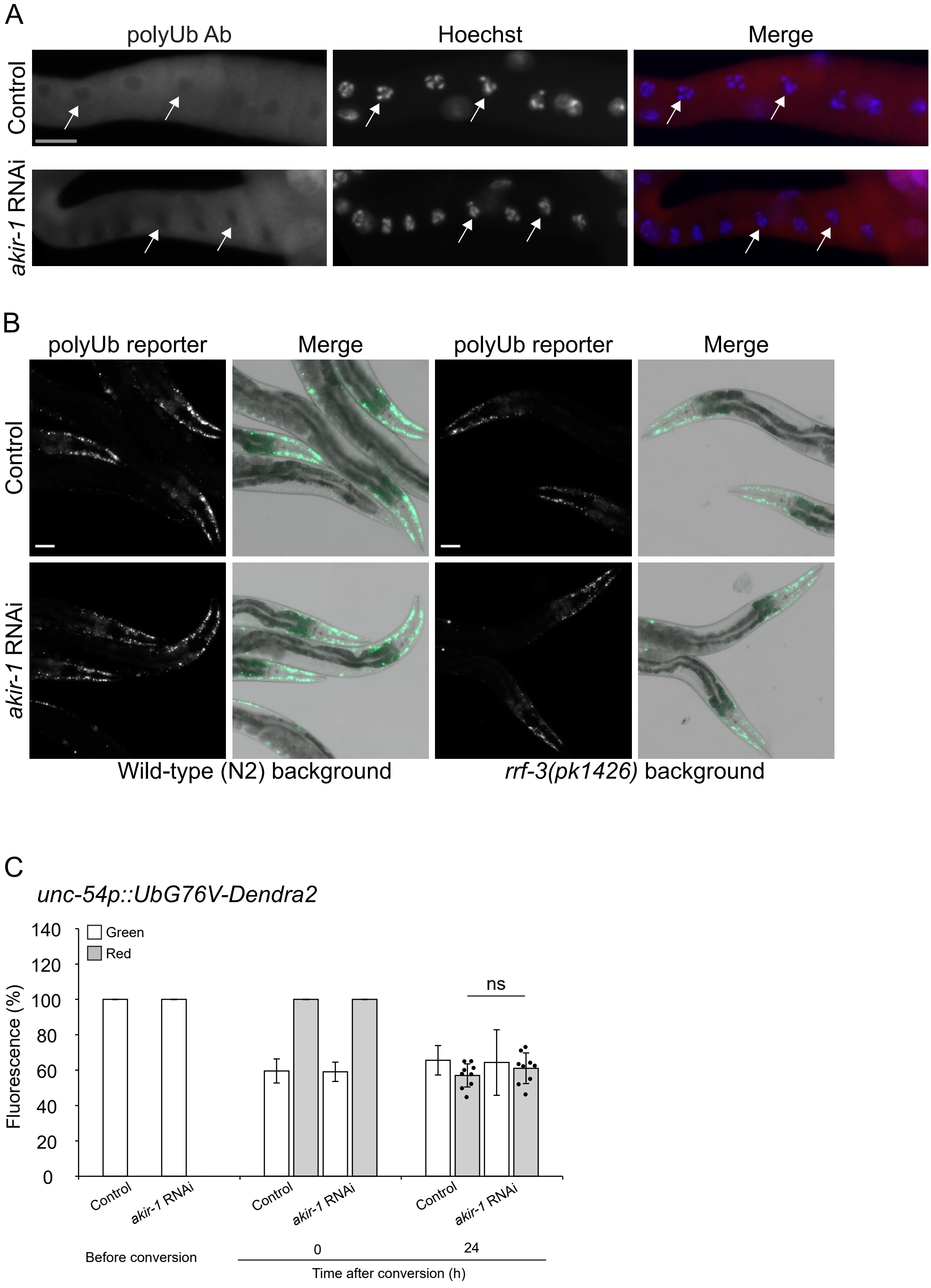


**Figure 7 — figure supplement 1.** Polyubiquitinated proteins do not accumulate in oocytes and body-wall muscle cells upon *akir-1* downregulation of $P_0$ generation. (**A**) Representative micrographs of polyubiquitin immunostaining (polyUb Ab) in dissected oocytes of control and *akir-1* RNAi–treated wild-type animals. Nuclei are visualized with Hoechst. Scale bar, 20 µm. White arrows point to representative nuclei. (**B**) Representative fluorescence micrographs of control and *akir-1* RNAi–treated animals expressing the polyubiquitin (polyUb) reporter (*unc-54p::UIM2-ZsProSensor)* in the body-wall muscle cells in N2 (two leftmost panels) or *rrf-3(pk1426)* (two rightmost panels) background. Merge represents overlay of fluorescence and bright-field images. Scale bars, 50 µm. (**C**) The graph shows the mean percentages of green and red fluorescence relative to the initial intensity (Before conversion) or at the indicated time points after conversion (0 h and 24 h) of control and *akir-1* RNAi–treated transgenic animals expressing photoconvertible UbG76V-Dendra2 (*unc-54p::UbG76V::Dendra2)* reporter proteins in the body-wall muscle cells. n= 3 independent experiments with triplicate images of 7 animals per image (total number of animals is 59 per treatment); Error bars, SD; ns, not significant.

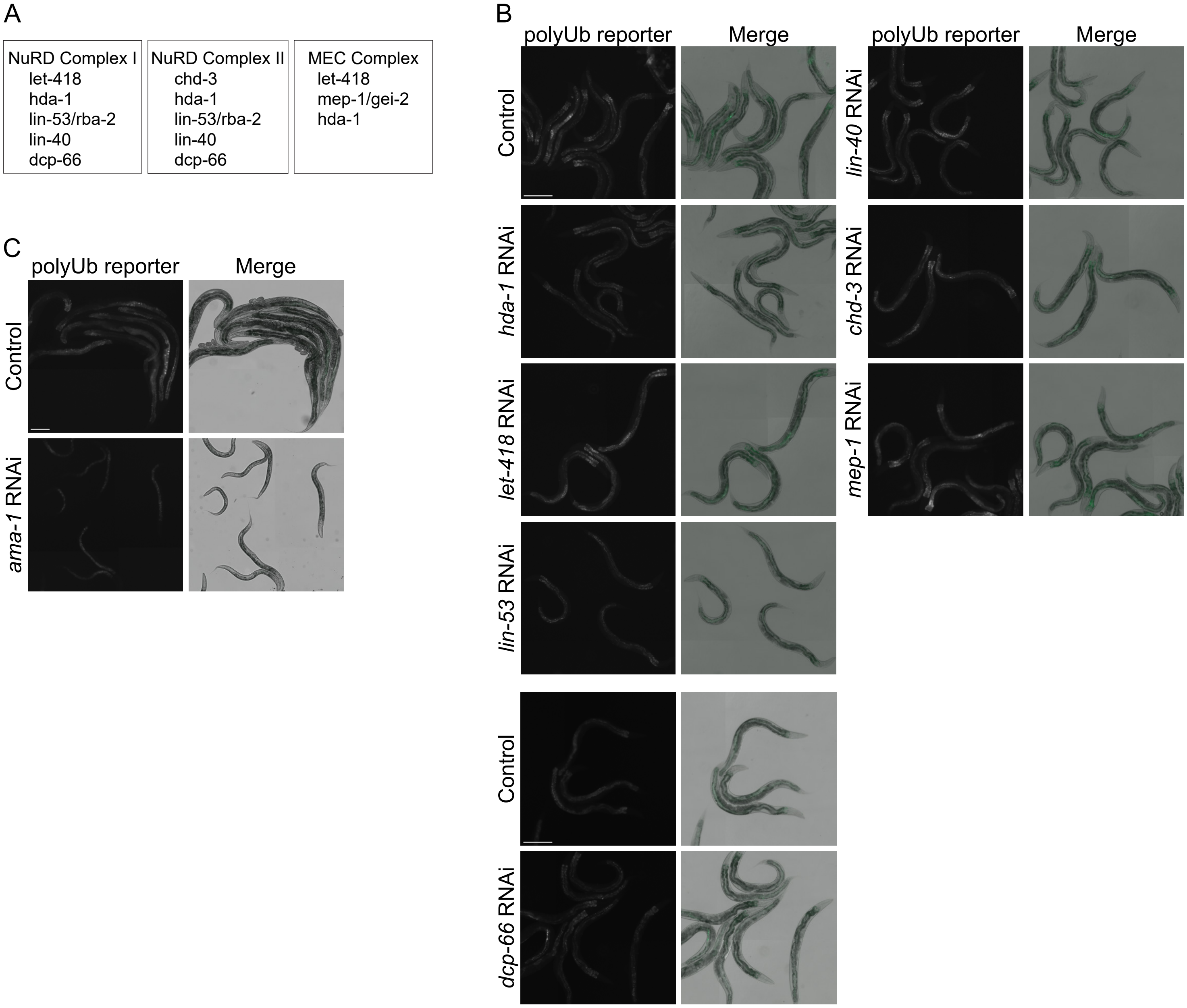


**Figure 8 — figure supplement 1.** Knockdown of genes of chromatin remodeling complexes or RNA polymerase II does not lead to nuclear accumulation of the polyubiquitin-binding reporter in intestinal cells. (**A**) The table shows genes of the *C. elegans* NuRD complex I and II, and the MEC complex. (**B, C**) Representative fluorescence micrographs of N2 animals expressing the polyubiquitin (polyUb) reporter (*vha-6p::UIM2-ZsProSensor)* in the intestinal cells and exposed to control or *hda-1*, *let-418*, *lin-53*, *lin-40*, *chd-3*, *mep-1, dcp-66* RNAi (**B**) (left panels), or *ama-1* RNAi (**C**) (left panels). Merge represents overlay of fluorescence and bright-field images (right panels). Scale bars, 200 μm.

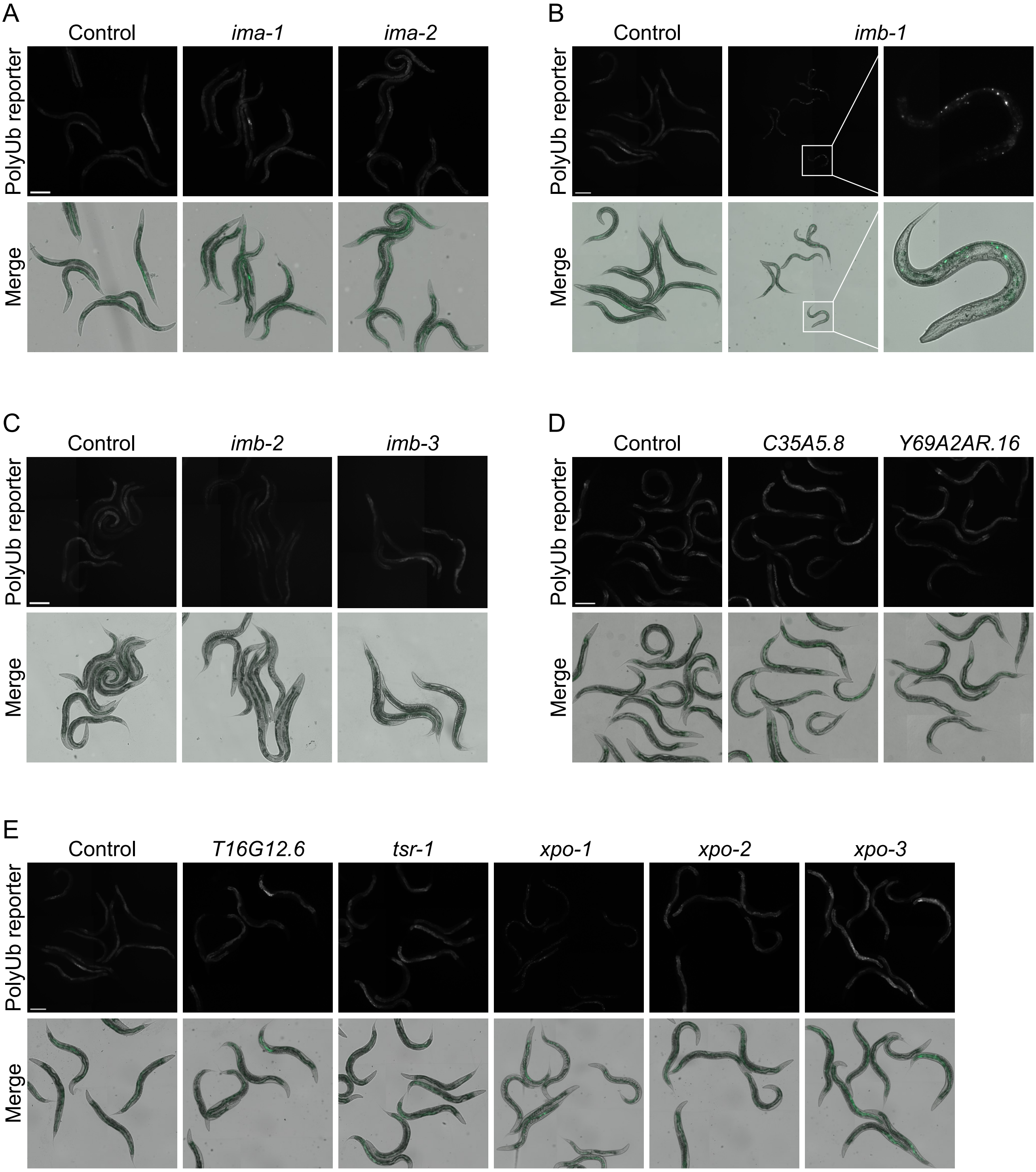


**Figure 9 — figure supplement 1.** Knockdown of *importin beta-1* leads to a partial nuclear accumulation of the polyubiquitin-binding reporter in intestinal cells. (**A-E**) Representative fluorescence micrographs of N2 animals expressing the polyubiquitin (polyUb) reporter (*vha-6p::UIM2-ZsProSensor*) in the intestinal cells and exposed to control or *ima-1*, *ima-2* (**A**), *imb-1* (rightmost panels show enlargements of the indicated areas) (**B**), *imb-2*, *imb-3* (**C**), *C35A5.8*, *Y69A2AR.16* (**D**), *T16G12.6*, *tsr-1*, *xpo-1*, *xpo-2*, or *xpo-3* RNAi treatment (**E**) (upper panels). Merge represents overlay of fluorescence and bright-field images (lower panels). Scale bars, 200 μm.
